# Endoplasmic reticulum stress protein GRP78 for ketamine’s antidepressant effects

**DOI:** 10.1101/2025.02.22.639685

**Authors:** Wanpeng Cui, Chen Shen, Wen-Cheng Xiong, Lin Mei

## Abstract

Ketamine is a fast-acting, long-lasting novel antidepressant. However, underpinning intracellular mechanisms remain unclear. We conducted an unbiased screening for genes that were less expressed in the prefrontal cortex (PFC) of mice that did not display antidepression-like effects to ketamine. GO analysis implicated endoplasmic reticulum and protein folding; in particular, GRP78, a stress-induced chaperone protein critical for protein folding, was reduced. We showed that GRP78 deficiency in PFC neurons induced depressive-like behaviors, whereas its overexpression produced anti-depression-like effects, revealing a novel function of GRP78. Prefrontal GRP78 was necessary for ketamine’s antidepressant-like effects. GRP78 was also required for ketamine to increase calcium activity and glutamatergic transmission. Enhancing GRP78 by viral infection and azoramide enabled non-responsive mice to respond to ketamine. Together, our results demonstrate that GRP78 is critical for ketamine to execute antidepressant effects by potentiating glutamatergic transmission in the PFC.

**Highlights:**

1. GRP78 was identified in non-biased screens for proteins that were lower in the PFC of mice that failed to exhibit antidepression-like effects after ketamine.
2. Reducing GRP78 in the PFC induced depressive-like behaviors while increasing GRP78 had opposite effects in wildtype mice.
3. GRP78 is required for ketamine to increase calcium activity in excitatory neurons and glutamatergic transmission.
4. Enhancing GRP78 by viral infection and by azoramide enabled non-responsive mice to respond to ketamine.

## Introduction

Major depressive disorder (MDD) is a widespread and debilitating psychiatric condition, affecting over 16% of the global population.^1^ Nevertheless, underpinning pathophysiological mechanisms remain poorly understood. First-line pharmacological treatments include selective serotonin reuptake inhibitors (SSRIs), serotonin-norepinephrine reuptake inhibitors (SNRIs), and dopamine-norepinephrine reuptake inhibitors (DNRI).^2–5^ The response and remission rates of these treatments remain suboptimal.^6–9^ Ketamine has recently emerged as a groundbreaking therapeutic due to its rapid and sustained antidepressant effects.^10,11^ At the neural circuit level, ketamine could enhance synchrony and dendritic spine density in PFC projection neurons, inhibition of PFC GABAergic interneurons and suppress burst firing in the habenula, a region implicated in aversive states.^12–15^ Molecular mechanisms include acting as a noncompetitive N-methyl-D-aspartate receptor (NMDAR) antagonist, enhancing the translation of brain-derived neurotrophic factor (BDNF), interaction with GluN2B in PFC interneurons and modulation of rapamycin (mTOR) signaling.^14,16–18^ Recent genetic insights highlight the Kcnq2 gene as a key modulator of its effects.^19^ Ketamine may undergo metabolism into bioactive metabolites.^20^ Despite these advances, the intracellular target and mechanisms underlying ketamine’s antidepressant efficacy remain incompletely understood.

The anti-depressive responses of ketamine are heterogenous among MDD patients, with ∼70% responding to ketamine treatment and ∼30% not responding.^11,21^ The response of geriatric patients was poor to repeat intravenous ketamine, although patients showed significant disassociated effects.^22^ Similar heterogenous effects of ketamine have been observed in mice. For example, ketamine could ameliorate depressive-like behaviors in ∼50% of susceptible mice after chronic social defeat stress (CSDS).^23^ However, the underlying mechanisms of ketamine response variety in animals remain poorly understood.

The endoplasmic reticulum (ER) has multiple functions including synthesis, folding, modification, and transport of proteins. It is also an organelle for protein quality control, relocating misfolded proteins into the cytosol for degradation by the ubiquitin-proteasome.^24^ The protein quality control system consists of glucose-regulated protein 78 (GRP78 also called HSPA5 or BiP), GRP94, chaperones, ATPases, and proteolytic systems.^25^ Accumulation of misfolded proteins could cause ER stress which could elicit adaptive responses to restore homeostasis; on the other hand, persistent ER stress could cause various diseases including neurodegenerative disorders. Recent studies have demonstrated chronic stress in life, a major risk factor for depression, can induce ER stress in the brain.^26,27^ ER stress markers such as GRP78, and Xbp1were reduced in patients with bipolar disorders, and associated with MDD.^28,29^ GRP78 is a stress-induced chaperone protein that belongs to heat shock protein family, critical for protein folding and assembly and export of misfolded proteins for degradation.^30^ In addition, GRP78 could regulate various signaling pathways, including the PI3K–AKT pathway and WNT–β-catenin signaling. Intriguingly, ketamine could increase the expression of ER stress-related proteins including GRP78.^31,32^ However, how GRP78 is involved in depression or ketamine’s effects remain unknown.

To identify molecular mechanisms that may be involved in mediating ketamine’s effects, we carried out a non-biased screen for molecules that were expressed at lower levels in mice that do not respond to ketamine. Our hypothesis was that MDD patients that fail to respond to ketamine may be due to the lack of a potential target protein(s). To test this hypothesis, a large cohort of mice were tested for their response to ketamine and categorized into mice that responded to ketamine and those that show no reduction in depressive-like behaviors (referred to as Res and NR mice, respectively).

RNA-seq analysis of PFC tissues revealed differentiated expressed genes (DEGs) that implicated protein folding and ER stress. In particular, GRP78 mRNA was downregulated in the PFC of NR mice. We showed that in wild type mice, GRP78 deficiency in PFC neurons induced depressive-like behaviors whereas its overexpression produced anti-depression-like effects, revealing a novel function of GRP78. Prefrontal GRP78 was necessary for ketamine’s antidepressant-like effects; enhancing the levels of GRP78 enabled NR mice to respond to ketamine. We have investigated cellular mechanisms of how GRP78 regulates depressive-like behaviors by analyzing calcium activity in free-moving mice, neuronal activity in brain slices and neuronal morphology. Together, our results demonstrate that GRP78 is critical to normal brain function and for ketamine to execute its antidepressant effects by potentiating glutamatergic transmission in the PFC.

## Results

### GRP78 downregulation in mice irresponsive to ketamine

We hypothesized that like humans, the responses to ketamine in mice may be heterogenous. To test this, C57Bl/6j mice were subjected to novelty suppressed feeding (NSF) and forced swimming (FST), behavior paradigms with high predictive validity to ketamine.^15,20,33,34^ In the NSF test, food-deprived mice were tested in a brightly lit arena containing a food pellet at the center and monitored for the feeding. Initially, mice were away from the pellet and after a period of exploration, began eating the food pellet. The time from the onset of the test to the time mice began feeding was defined as latency to feeding, which indicates depressive-like and anxiety levels. In agreement with previous reports,^15,20^ mice treated with ketamine (10 mg/kg, i.p., 24 hours) displayed an overall reduction in the latency to feeding in NSF, compared with mice treated with vehicle (Veh) (Veh, 333.9 ± 183.3 s vs ketamine, 220.6 ± 195.9 s, p = 0.014), demonstrating antidepression-like effects of ketamine (Figure 1A). To identify mice that were unable to respond to ketamine, a large cohort of naive mice (n = 109) were treated with Veh and subjected to NSF at day 2 (NSF1) and treated with ketamine at day 7 and tested at day 8, recorded for latency to feeding in NSF (NSF2) (Figure 1B). Notice that the two NSF tests were separated by 7 days and in different beddings, to eliminate pre-exposure effect. As shown in Figure S1A-S1B, latency to feeding was similar between NSF1 and NSF2 treated with saline, suggesting limited pre-exposure effects in our model. The overall population of 109 mice reduced latency after ketamine treatment (Veh, 278.4 ± 128.7 s vs ketamine, 208.7.6 ± 170.6 s, p = 0.023, Figure 1C). To determine the responsiveness of each mouse, the change in latency to feeding of individual animals (i.e., Δlatency) was calculated by subtracting the latency of NSF1 from that of NSF2. A negative value of Δlatency would indicate a reduction in latency to feeding by ketamine (see equation1). As shown in Figure 1D, the majority (66 of 109 naïve mice, 60.5%) responded to ketamine with Δlatency values smaller than zero. In contrast, 43 mice (39.5%) display Δlatency values larger than zero, suggesting that these mice failed to reduce the latency to feeding, suggesting that they were unable respond to ketamine (Figure 1E).

**Figure 1.**
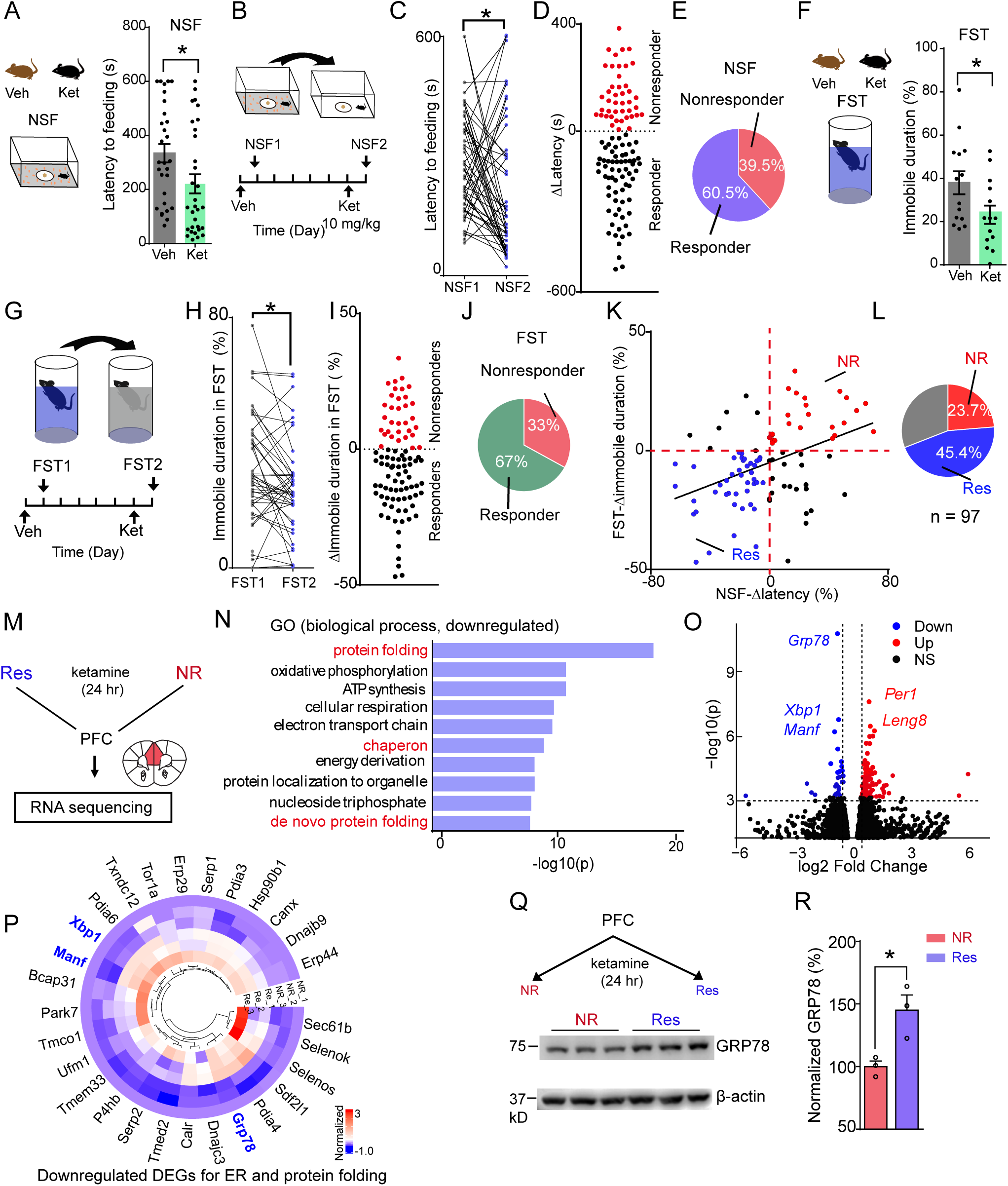
GRP78 downregulation in mice irresponsive to ketamine. (A) Ketamine reduces the latency to feeding in NSF. Two groups of mice were injected with ketamine (Ket, 10 mg/kg) or Veh 24 hours before tests. Mann Whitney test, p = 0.014, U = 284.0, n = 29-31 mice for each group. (B-C) Test of same group of mice in two consecutive NSF tests. (B) Tests were performed with 7 days apart, with different bedding in the box (grey and white on floor). (C) Ketamine reduced latency to feeding in the two NSF. p = 0.023, W = −470.0, n = 50 mice, Wilcoxon rank test. (D) Δlatency in NSF before and after ketamine. Red dots, nonresponders, black, responders. n = 109 mice. (E) Percentage of nonresponders and responders from D. (F) Reduction of immobile duration in FST in two group of mice injected with ketamine or Veh. T-test, p = 0.039, t = 2.19. n = 14 mice for each group. (G-H) Same mice tested in two FST tests. (G) Schematic diagram of two FST, with transparent or covered tank. (H) Reduction of immobile duration after ketamine. Paired t-test, t = 2.65, p = 0.011, n = 44 mice. (I) Δimmobile duration in FST before and after ketamine. Red dots, nonresponders, black, responders. (J) Percentage from I. n = 97 mice. (K-L) Identification of NR and Res mice by Δlatency and Δimmobile duration. (K) Correlation analysis of Δlatency in NSF (normalized) and Δimmobile duration in FST. n = 97 mice. p = 0.0002, r = 0.369. Pearson r analysis. Red dash lines, threshold was set to 0. Red dots, NR; blue, Res mice. Black dots represent single responders or non-responders in either NSF or FST. (L) Percentage of NR and Res mice from K. (M) Schematic showing the RNA sequencing using the PFCs from both Res and NR mice. Mice were injected with ketamine 24 hours before sampling. (N) Gene ontology analysis using biological process. Red, protein folding related pathways. (O) Volcano plotting of the DEGs. Red dots, upregulated genes, blue, downregulated genes, black, not significant. Dash line, threshold for significance. (P) Downregulated genes for ER and protein folding. Scale,[-1 3]. (Q-R) Protein of GRP78 was reduced in NR mice compared to Res after ketamine in PFC. (Q) Western blotting of GRP78 in PFC of Res and NR mice. Ketamine (10 mg/kg, i.p.) was administrated 24 hours before sampling. β-actin was used as internal control. (R) Bar graph of Q. T-test, t = 3.49, p = 0.025. n = 3 mice for each group. * p < 0.05. Data, mean ± SEM.

To determine this inability was detected in a different test, mice were subjected to FST, mice were monitored struggling and immobile behaviors, where immobile duration server as an indicator of depressive-like level.^35^ Ketamine-treated mice reduced in immobile duration, compared with Veh-treated mice (Veh, 37.9 ± 19.9% vs ketamine, 23.1 ± 15.9%, p = 0.039) (Figure 1F), in agreement with previous reports.^15,20,34^ To determine whether the ketamine response was heterogenous in FST, the same cohort of mice were subjected to two FST tests by using a similar strategy as described above (Figure 1G). Preliminary results indicate that immobile duration of two FSTs were similar under two saline injections, suggesting no pre-exposure effects (Figure S1C-S1D). However, the immobile durations were reduced after ketamine treatment (Veh, 32.9 ± 17.9% vs ketamine, 27.1 ± 15.7%, p = 0.011), indicative of ketamine’s antidepressant effects (Figure 1H). Individually, however, ketamine-induced reduction in immobile duration was observed in 65 of 97 naïve mice (67%), Δimmobile duration values smaller than zero (black dots in Figure 1I). In contrast, 32 mice (33%) display Δimmobile duration larger than zero (red dots in Figure I), suggesting that these mice failed to respond to ketamine (Figure 1J).

Notably, the Δlatency positively correlated with that Δimmobile duration in FST (Figure 1K, p = 0.0001, n = 97 mice). Of total 97 mice, 23.7% were nonresponders and 45.4% responded in both NSF and FST (red and blue dots, respectively in Figure 1K-1L). By using two different behavioral paradigms, we demonstrate that C57Bl/6j mice displayed heterogenous responses to ketamine to alleviate depressive-like behaviors. In so doing, we were able to categorize mice into mice that respond to ketamine (referred to as Res mice) and those that failed to respond to ketamine (referred to as NR mice, Figure 1L). Notice that the responsiveness of NR as well as Res mice was similar to ketamine at two repeated administration of ketamine (Figure S2A-S2E) or a higher dose observed similar population of NR and Res mice (20 mg/kg, NR, 26.7% and Res, 53.3%) (Figure S3A-S3H). In the absence of ketamine, Res and NR mice showed no difference in latency to feeding in NSF or immobile duration in FST (Figure S4A-S4E), excluding potential differences of tested mice in innate depressive-like behaviors.

To validate whether such categorized mice would respond in a similar way after stress, we subjected NR and Res mice to chronic social stress (CSDS), a paradigm that is known to cause depressive-like behaviors (Figure S1E).

Earlier reports indicate that ketamine alleviate depressive-like behaviors in social avoidance (SA) in CSDS-stressed susceptible mice.^20,23^ Susceptible NR mice showed no improvement after ketamine treatment in SA (Veh, 37.2 ± 33.9% vs ketamine, 46.0 ± 26.1%, p = 0.731, Figure S1F). As control, CSDS-stressed susceptible Res mice increased social interaction after ketamine treatment (Veh, 54.7 ± 16.5% vs ketamine, 92.9 ± 35.9%, p = 0.023, Figure S1G). These results revealed that ketamine had little effect on NR mice that were nonresponsive to ketamine for CSDS-induced depressive-like behaviors, demonstrating lacking antidepression-like effects. Taken together, we characterize the responses of WT animals to ketamine in both the NSF and FST, and validate these findings in the CSDS, indicating heterogeneous response to ketamine.

To investigate mechanisms of NR mice’s inability to respond to ketamine, we identified differentially expressed genes (DEGs) in the PFC, a target brain region that is implicated in depression and regulation by ketamine,^36,37^ between NR and Res mice (Figure 1M). Among 2486 DEGs (p < 0.05), 1375 genes were downregulated (Table 1), whereas 1111 genes increased in NR mice (Table 2), compared with Res mice (Figure S5A-S5B).

Gene ontology (GO) analyses of down-regulated DEGs revealed several biological processes related to ER stress, and protein folding was most significant(p = 2.83E-16, Figure 1N). Among 1375 down-regulated DEGs, 180 genes (15.2%) were ER proteins, and 28 genes for ER and protein folding (Figure 1P, Table 1). In addition, up-regulated DEGs were implicated extracellular matrix and ligand-gated calcium channels (Figure S5C). Of down-regulated DEGs was glucose-regulated protein 78 (GRP78) (p =1.25E-11) that showed most significance in statistical analysis (Figure 1O). GRP78 is a 78-kDa ER-resident molecular chaperone and a critical effector in ER stress to correct and clear misfolded proteins.^38,39^ To validate this finding by RNA sequencing, we performed western blotting analysis. As shown in Figure 1Q-1R, ketamine increased GRP78 in PFC of Res mice after ketamine, compared with that of NR mice, consistent with the RNA-seq data. Notice that GRP78 level was not different between NR and Res without ketamine injection, consistent with similar behaviors in baseline (Figure S5D-S5E). Ketamine is an antagonist of NMDAR, but its mRNA or protein was similar between NR and Res mice (Figure S5D and S5F). However, GRP78 levels were increased in human neuroblastoma SH-SY5Y cells by ketamine, suggesting that ketamine was able to induce GRP78 (Figure S5G-S5H). Taken together, these results demonstrate an association of lower levels of GRP78 with the inability of NR mice to respond to ketamine and suggest the involvement of ER stress and protein folding may be involved in depression and its response to ketamine.

### Neuronal GRP78 for ketamine’s antidepressant effects in naïve and stressed mice

Lower levels of GRP78 in NR mice suggest that GRP78 may be necessary for mice to respond to ketamine. Immunostaining indicated that GRP78 was enriched in PFC neurons (green in Figure 2A). To investigate its role in depressive-like behaviors, we injected AAV-hSyn-Cre-GFP or AAV-hSyn-GFP (as control) in the PFC of *Grp78(f/w)* mice whose Exon5-Exon7 were flanked by LoxP cassettes(Figure S6A and Figure 2B).^40^ Viral injection led to the expression of GFP in the PFC (green in Figure 2C). In mice injected with Cre, the staining of anti-GRP78 antibody was reduced in the PFC (purple in Figure 2C). Western blotting indicated ∼40% reduction in GRP78 in viral Cre-injected PFCs, compared with controls (Figure 2D-2E). Intriguingly, Cre virus-injected mice displayed increased latency to feeding in NSF, compared with mice injected with GFP virus (latency for feeding: Cre virus, 397.2 ± 160.3 s vs GFP virus, 205.6 ± 86.9 s, p = 0.047, Figure 2F). In accord, in FST, mice injected with Cre virus had higher immobile duration, compared with mice injected with control virus (Cre, 52.3 ± 19.9% vs GFP, 24.4 ± 10.4%, p = 0.0001) (Figure 2G). These results indicates that loss of GRP78 in PFC might induce depression-like behaviors.

**Figure 2.**
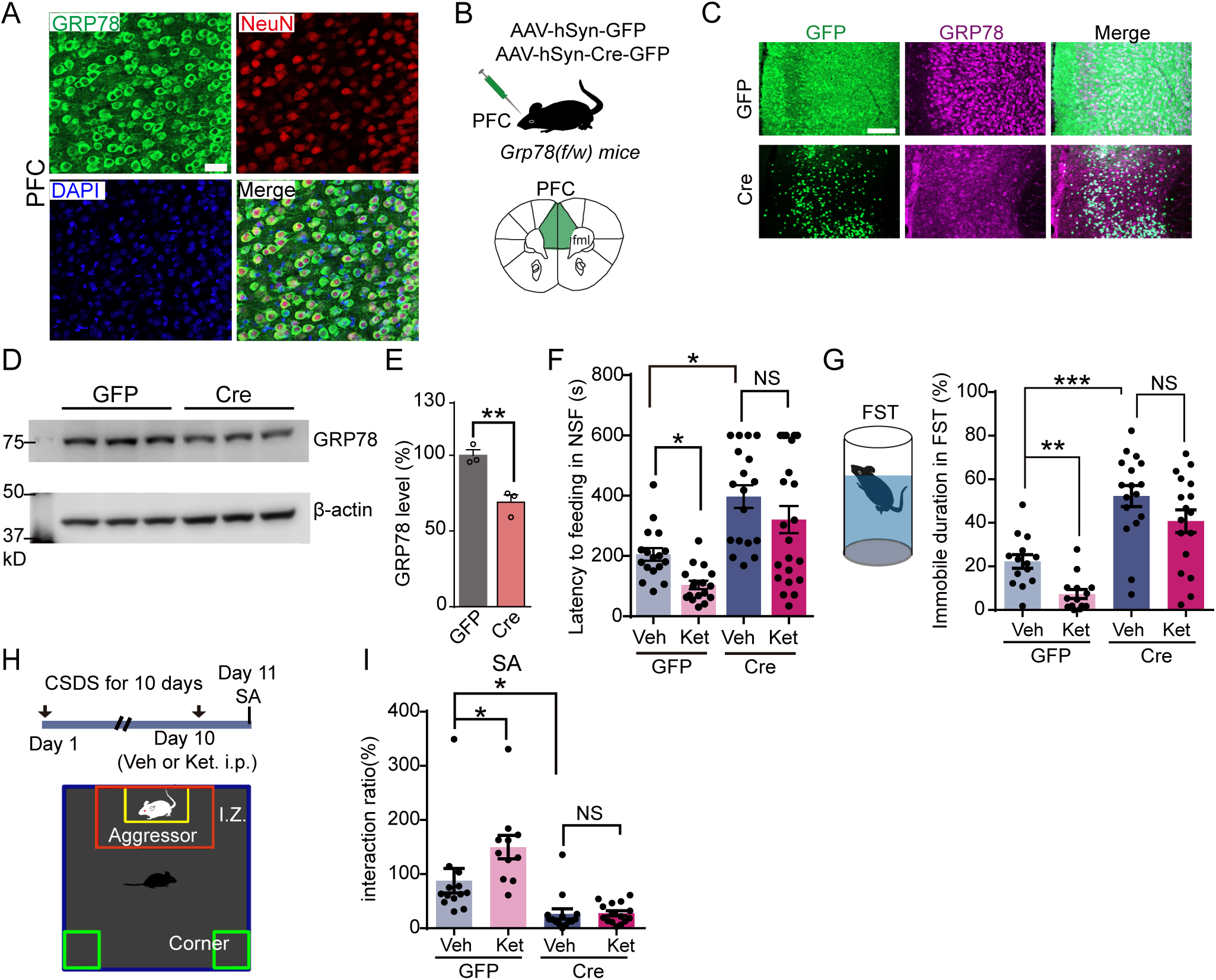
Neuronal GRP78 for ketamine’s antidepressant effects in naïve and stressed mice. (A) Enrichment of GRP78 in PFC neurons. Antibody staining of GRP78 (green) and NeuN (red) in coronal sections of PFC. Blue, DAPI, scale, 50 µm. (B-C) Reduction of GRP78 in PFC after AAV. (B) Schematic diagram showing AAV injection in bilateral PFC of Grp78(f/w) mice. (C) Expression of GRP78 in PFC after AAVs injection. GFP, green, GRP78, purple. Scale, 100 µm. (D-E) Reduced GRP78 expression in PFC. (D) Western blotting of GRP78 in PFC of GFP and Cre injected mice. β-actin was used as internal control. (E) Quantification in C. p = 0.0076, t = 4.99. T-test. n = 3 mice for each group. (F) Reducing GRP78 abolished ketamine’s effects in NSF. K = 31.1. Kruskal-Wallis test, followed by Dunn’s multiple comparison. n = 17-22 mice for each group. (G) Immobile duration in FST. F_(3, 58)_ = 31.6. n = 14-17 mice for each group. (H-I) SA test after CSDS. (H) CSDS and schematic diagram of SA. GFP or Cre injected mice were treated with CSDS for 10 days before SA. Red rectangle, interaction zone (I.Z.). (I) Interaction ration in SA. F_(3, 49)_ = 13.6. One-way ANOVA, followed by Tukey’s *post hoc*. n = 11-15 mice for each group. Data, mean ± SEM, * p < 0.05, ** p< 0.01, ***p < 0.001. NS, not significant.

To determine whether GRP78 deficiency altered the response to ketamine, mice injected with GFP virus were subjected to NSF tests. As shown in Figure 2F ketamine reduced the latency to feeding in GFP group (205.6 ± 86.9 s with Veh vs 103.8 ± 57.4 s with ketamine, p = 0.044), in agreement with previous reports.^20^ Remarkably, these effects were not observed in mice injected with Cre virus (latency to feeding: 397.2 ± 160.3 s with Veh vs 320.7 ± 210.7 s with ketamine, p = 0.62). In the FST, ketamine injection reduced the immobile duration from 24.4 ± 10.4% to 7.3 ± 7.90% (p = 0.04) in mice injected with GFP virus (Figure 2G). In contrast, no difference was observed in immobile duration of Cre virus-injected mice treated with Veh and ketamine (52.3 ± 19.9% with Veh vs 40.8 ± 21.5% with ketamine, p = 0.187, Figure 2G). These data demonstrate that GRP78 deficiency prevented mice from responding to ketamine, suggesting a necessary role of GRP78 in executing ketamine’s anti-depressive like effect.

Next, we determined whether GRP78 is required for stressed mice to respond to ketamine. Mice were injected with GFP and Cre viruses, respectively and subjected to CSDS, followed by SA, a most responsive paradigm to reveal depressive-like state to CSDS (Figure 2H).^41^ Social interaction ratio was reduced in Cre virus-injected mice, compared with GFP virus-injected mice treated with Veh (26.5 ± 34.6% with Cre virus 87.8 ± 81.9% with GFP virus, p = 0.031) (Figure 2I), indicating that GRP78 deficiency may be causal to depressive-like behaviors, in agreement with findings in Figure 2F-2G. In CSDS-stressed, GFP virus-injected mice, ketamine treatment increased social interaction ratio (87.8 ± 81.9% with Veh vs 149.8 ± 71.5% with ketamine, p = 0.044), indicative of antidepression-like effects (Figure 2I). In contrast, the social interaction ratio in CSDS-stressed, Cre virus-injected mice remained low after ketamine treatment and showed no difference from that of Veh injected mice (26.5 ± 34.6% with Veh vs 27.8 ± 18.5% with ketamine, p > 0.99) (Figure 2I). These results demonstrated that prefrontal GRP78 was necessary for ketamine’s antidepressant-like effects, identifying a potential target of ketamine.

### Res mice became non-responsive by reducing GRP78 levels

We showed that like humans,^11,21^ mice were heterogenous in responding to ketamine and could be categorized into Res and NR groups that were associated with GRP78 levels (Figure 1O-1R). Having demonstrated a necessary role of GRP78 in mediating ketamine’s antidepressant-like effects, it would be interesting to determine whether manipulating GRP78 levels in PFC could alter the responses in Res and NR mice. To this end, Grp78f/w mice were screened for those that were responsive and irresponsive to ketamine (i.e., Res and NR mice, respectively). First, we tested whether reducing GRP78 levels alters the response of Res mice. To this end, Res mice were screened and divided into two groups that were injected with GFP or Cre virus, respectively (Figure 3A). As shown in Figure S6B-S6C, GRP78 level was reduced by ∼45% in Res mice with Cre compared to GFP injection. 24 days later, mice were treated with ketamine, followed by NSF (NSF3 in Figure 3A). Notice that the second dose of ketamine was given ∼25 days after the screening. In preliminary experiments, ketamine given ∼25 days apart was able to produce similar antidepressant effects in mice (Figure S2A-S2D). As shown in Figure 3B, Res mice remained responsive to ketamine after injection with GFP in NSF (latency: 319.0 ± 192.1 s with Veh, compared to 202.5 ± 192.9 s, p = 0.005 with ketamine, and 137.1 ± 132.4 s, p = 0.001, with GFP+ketamine), in agreement with previous data (Figure S2A-S2B). However, ketamine was not as effective in Res mice injected with Cre virus, with increased latency (172.4 ± 143.8 s before Cre vs 337.7 ± 201.3 s, p = 0.043 after Cre, Figure 3C). As shown in Figure 3D-3E, 60% of Res mice (red dots in 3D, Δlatency ≥ 0) injected with Cre virus failed response to ketamine (Δlatency: GFP group, −81.9 ± 168.8 s vs 4.0 ± 203.3 s in Cre, p = 0.025). As control, almost all GFP virus-injected Res reacted to ketamine with anti-depressive-like responses (100%, Δlatency < 0, blue dots in Figure 3D), suggesting that the effect of Cre virus was not mediated by AAV infection alone.

**Figure 3.**
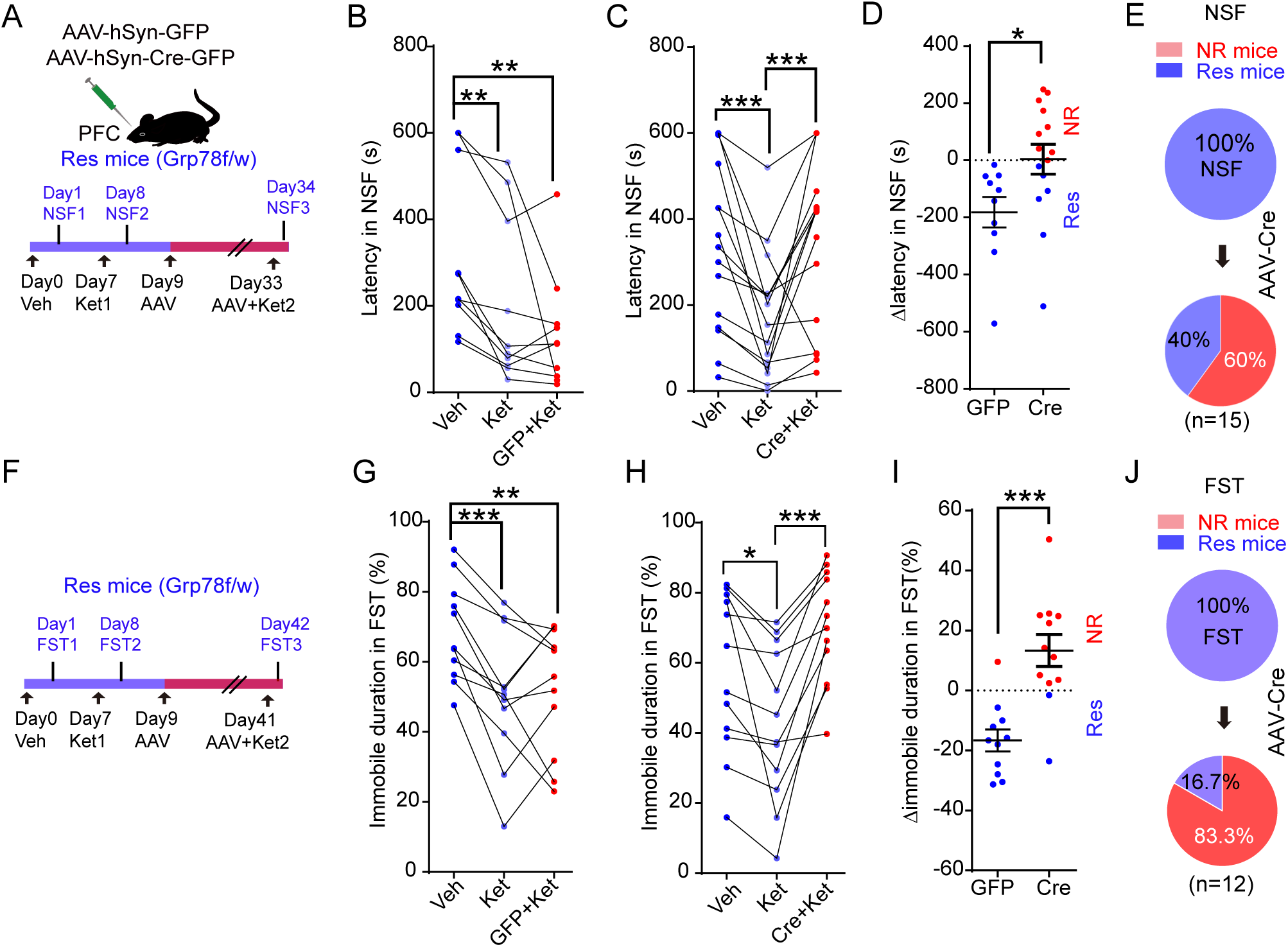
Res mice became non-responsive to ketamine by reducing GRP78 levels. (A-E) Res mice reduced responses to ketamine after AAV in NSF. (A) AAVs injection and experimental design. Grp78(f/w) mice were screened for Res mice before AAV injections in PFC. (B) Latency in NSF after GFP, Friedman’s F = 15.2. n = 10 mice. (C) Latency in NSF after Cre, Friedman’s F = 19.9, n = 15 mice. Friedman test, followed by Dunn’s *post hoc*. (D) Δlatency calculated from B and C (NSF3-NSF1). Blue, Δlatency < 0, Res mice. Red, Δlatency ≥ 0, NR mice. t = 2.39, t-test. (E) Pie chart of population after Cre based on Δlatency in Figure 3D. n = 15 mice. (F-J) FST. (F) Experimental design. (G) Immobile duration in GFP group. F_(10, 20)_ = 8.38. n = 11 mice. (H) Cre. F_(11, 22)_ = 6.28. n = 12 mice. Repeated measures ANOVA, followed by Tukey’s *post hoc*. (I) Δimmobile duration calculated from G and H (FST3-FST1). T-test, t = 4.55. (J) Percentage of Res or NR in FST from Cre injection group from Figure 3I. * p < 0.05, ** p < 0.01, ***p < 0.001. Data, mean ± SEM.

Together, the above results demonstrate that reducing GRP78 levels in the PFC reduced the responses of Res mice, revealing again a critical role of GRP78. This notion was supported by results from FST studies. As shown in Figure 3F-3J, as a population, ketamine elicited antidepressant-like response in Res injected with GFP virus (68.4 ± 14.2% with Veh, to 50.3 ± 19.2% with ketamine, p = 0.0005, and to 52.0 ± 17.9% with GFP+ketamine, p = 0.003, Figure 3G). However, those injected with Cre virus failed to display the antidepression-like effects of ketamine, rather increased the immobile duration (42.8 ± 22.1%, p = 0.035 with ketamine, compared to 70.4 ± 15.9% with Cre+ketamine, p = 0.0004, Figure 3H). The Δimmobile duration was increased in Cre injected mice compared with GFP control, suggesting pro-depression-like effects (p = 0.0002, Figure 3I). 10 out of 12 (83.3%, Δimmobile duration ≥ 0) Res mice after Cre-virus injection became non-responsive in FST (Figure 3J).

### Anti-depressive-like effects of enhancing GRP78 levels

Having demonstrated a necessary role of GRP78 in ketamine’s antidepressant effects, we next determined whether increasing GRP78 levels would be sufficient in controlling the responsiveness to ketamine. To this end, GRP78-P2A-mCherry AAV was generated (Figure 4A); its injection increased GRP78 levels in PFC, compared with that with AAV-mCherry (green in Figure 4B, and Figure S7A-S7B). Intriguingly, GRP78 virus-injected mice were less in the latency to feeding in NSF, compared with mice injected with mCherry virus (Figure 4C). They also reduced immobile duration in FST (Figure 4D). These results indicate that mice with elevated GRP78 levels in the PFC showed less depressive-like behaviors in NSF and FST. Similar results were obtained with stressed mice; as depicted in Figure 4E-4F, GRP78 virus-injected CSDS-stressed mice displayed more interaction with the unfamiliar aggressors, compared with control mice (i.e., mCherry virus-injected CSDS-stressed mice).

**Figure 4.**
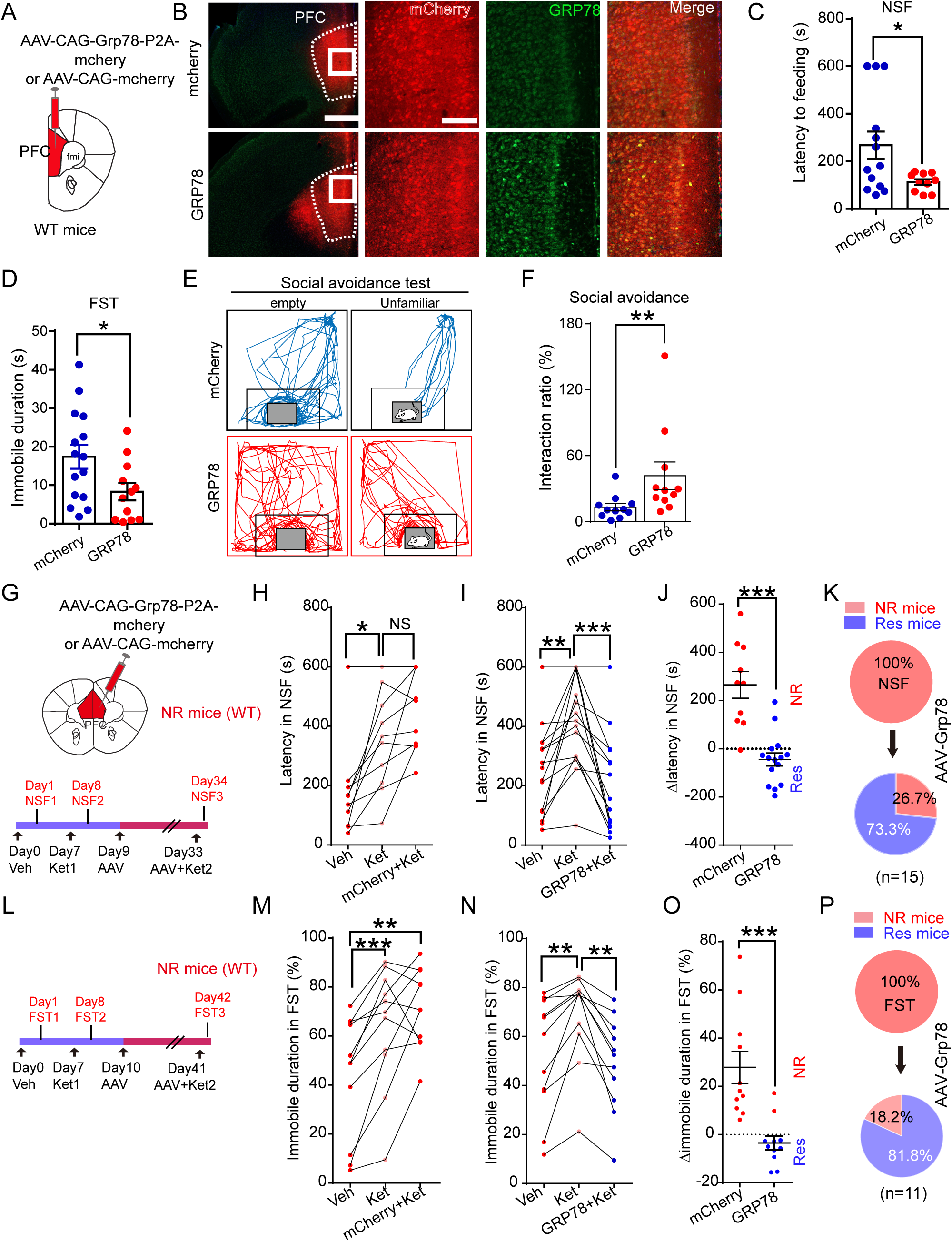
Anti-depressive-like effects of enhancing GRP78 levels. (A) Schematic diagram showing injection of AAVs in WT mice. AAV9 harboring mCherry or Grp78-mCherry were injected into the bilateral PFC. (B) Coronal brain sections showing expression of GRP78 (green) and mCherry (red) in PFC. Scale bar, 500 µm and 200 µm. (C-F) increasing GRP78 in the PFC produced anti-depression-like effects. (C) Latency to feeding in NSF. p = 0.049, U = 33.0. Mann-Whitney test. n = 10-13 mice for each group. (D) FST. p = 0.032, t = 2.27. n = 12-15 mice for each group. (E-F) SA after CSDS. (E) Traces of animals in SA after CSDS. Blue, mCherry, red, GRP78. Rectangle, I.Z.. (F) Interaction ratio in SA. p = 0.038, t = 2.23. n = 11 mice for each group. T-test. (G-K) Increasing GRP78 in PFC of NR mice increased response to ketamine in NSF. (G) AAV injections in PFC of NR mice and experimental design. WT mice were screened for NR mice using NSF and FST, then for AAV injection. (H) mCherry injected mice in NSF1 (Veh), NSF2 (Ket) and NSF3 (mCherry+Ket). Friedman’s F = 14.9, n = 10 mice. (I) Latency to feeding in GRP78 group. Friedman’s F = 23.3, n = 15 mice. Friedman test, followed by Dunn’s *post hoc*. (J) Δlatency calculated from H and I. t-test, t = 5.52. (K) Pie chart of Figure 4J (GRP78 group). n = 15 mice. (L-P) NR mice increased response to ketamine after AAV-GRP78 in FST. (L)Time course of experiments. (M) Immobile duration in mCherry injected mice in FST. F_(10, 20)_ = 6.68, n = 11 mice. (N) Immobile duration in GRP injected mice. F_(10, 20)_ = 15.3, n = 11 mice. Repeated measures ANOVA, followed by Tukey’s *post hoc*. (O) Δimmobile duration (FST3-FST1) calculated from M and N. t = 4.30. t-test. n = 11 mice for each group. (P) Pie chart of Figure 4O from GRP78 group. n = 11 mice. NS, not significant, ** p < 0.01, ***p < 0.001. Data, mean ± SEM.

These data suggest that elevating GRP78 levels in PFC produced anti-depression-like effects in naïve as well as stressed mice, in support of our hypothesis. GRP78 was identified as a gene whose mRNA and protein were reduced in the PFC of mice that were unable to respond to ketamine (i.e., NR mice), compared with Res mice (Figure 1O-1R). Results of our loss-of-function studies support the hypothesis that GRP78 is required for ketamine’s antidepressant effects (Figure 2 and Figure 3). This hypothesis predicts that increasing GRP78 levels in the PFC may enable NR mice to respond to ketamine with antidepressant effects. To test this, we took the same approach (as illustrated in Figure 3) by subjecting naïve mice to ketamine treatment and based on their response to ketamine, categorized them into NR and Res groups. NR mice were injected with mCherry or GRP78 viruses, respectively and ∼3 weeks later, tested for ketamine’s antidepressant effects (Figure 4G and Figure 4L). As described earlier, the second dose of ketamine given ∼25 days after the first was as effective as the first dose (Figure S2). Ketamine remained ineffective in mice injected with mCherry virus as the latency to feeding and immobile duration were higher in ketamine-treated group in NSF (175.8 ± 160.3 s with Veh to 441.5 ± 131.7 s with ketamine and mCherry, p = 0.001, Figure 4H) and in FST (43.7 ± 24.9% with Veh to 71.6 ± 16.1% with ketamine and mCherry, p = 0.005, Figure 4M). Remarkably, GRP78 virus-injected NR mice reduced the latency to feeding in NSF (415.1 ± 155.6 s with ketamine, to 205.5 ± 164.1 s with GRP78+ketamine, p = 0.002, Figure 4I), and decreased the immobile time in FST (68.5 ± 18.9% with ketamine to 48.9 ± 19.3% with GRP78+ketamine, p = 0.0013, Figure 4N). In accord, Δlatency and Δimmobile duration values were reduced by GRP78 (Δlatency: 265.2 ± 175.7 s in mCherry vs −44.6 ± 105.6 s in GRP78, p < 0.0001; for Δimmobile duration: 27.9 ± 22.2% with mCherry to −3.5 ± 9.7% with GRP78, p = 0.0004) (Figure 4J and 4O). Overall, after GRP78 viral injection, 73.3% of NR mice became responsive to ketamine in NSF, and 81.8% in FST (Figure 4K and 4P). These results demonstrate that increasing GRP78 levels in the PFC enable NR mice to respond to ketamine with antidepressant effects.

### Impaired glutamatergic transmission in NR mice

Having demonstrated a critical role of GRP78 in controlling ketamine’s responsiveness, we sought to investigate underpinning cellular mechanisms. Ketamine was known to promote the neuronal activity in the PFC that is implicated in its antidepressant effects.^42^ Therefore, we determined in free-moving mice whether PFC neuronal activities were different between NR and Res mice and whether the activities may be altered by ketamine. To this end, Nex-Cre mice were injected into the right side of PFC with AAV9-hSyn-Flex-GCaMP6f and 3 weeks later, implanted with a miniaturized integrated fluorescence microscope into the right PFC (Figure 5A-5B). Mice were then screen for ketamine’s response for classification into NR and Res groups, which were subjected to NSF. Calcium signals of excitatory neurons (ExNs) in the FOV (field of view) were synchronized with NSF behaviors by a data acquisition system (DAQ) (Figure 5C). LED power was below 0.3 mW/mm^2^to minimalize the photobleaching (see methods for details). As shown in Figure 5B and Figure 5D, a significant number of PFC ExNs were visualized.

**Figure 5.**
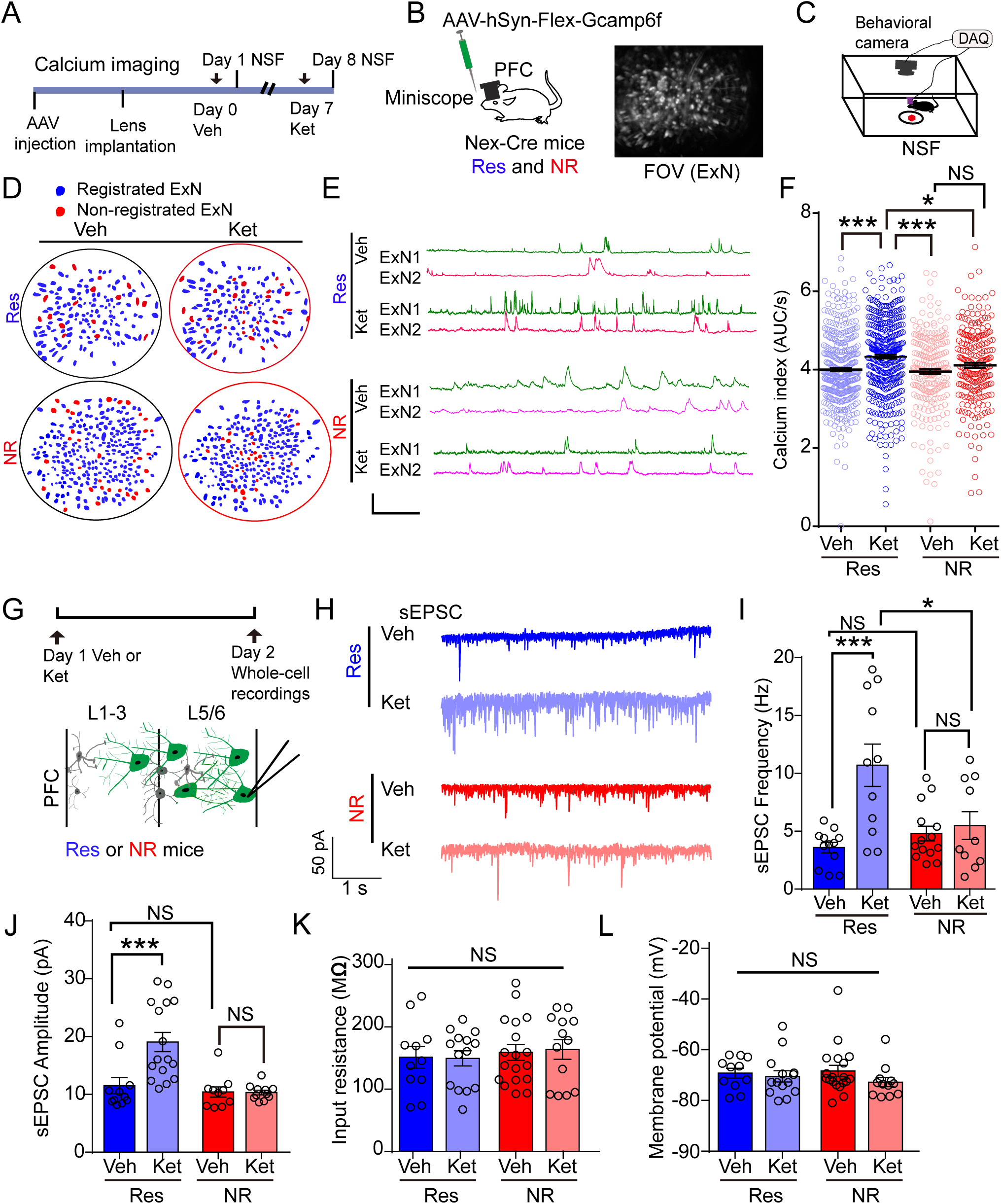
Impaired glutamatergic transmission in NR mice. (A-C) Calcium imaging of PFC excitatory neurons in Res and NR mice. (A) Experimental design. (B) AAV injection and representative field of view (FOV) of calcium signal from excitatory neurons in PFC. Nex-Cre mice (including Res and NR) were injected with AAV-hSyn-Flex-GCaMP6f in PFC. Ketamine was injected ∼ 24 hours before recordings. (C) Schematic diagram showing synchronization of behavior and calcium activity in NSF. Behaviors and calcium signal were recorded by a camera or miniscope. Red, food pellet. (D) Representative spatial footprints showing registration of ExNs in Veh and ketamine administration. Blue cells, registered neurons, red, non-registered ExNs. (E) Representative traces showing the calcium activity before and after ketamine in NSF. Calcium activity of two representative EXNs (color lines) under saline or ketamine in NSF. Red shades represent feeding trials. Scale, 10% ΔF/F, 100 s. (F) Ketamine potentiated calcium activity in feeding stages of NSF. Scatter plot showing the quantification of calcium index from registration neurons in feeding periods, and under the effects of Veh or ketamine. Res (Veh vs Ket), p < 0.0001; NR (Veh vs Ket), p = 0.263, Res+Veh vs NR+Veh, p = 0.92, Res+Ket vs NR+Ket, p = 0.026. F_(3,1206)_ = 11.5. Bars, mean ± SEM. n = 225-407 neurons from 3 mice of each group. One-way ANOVA, followed by Tukey’s post hoc. (G) Schematic diagram of whole-cell recordings in PFC layer 5 pyramidal neurons in Res and NR mice. Veh (saline) or Ket (20 mg/kg) was injected to Res and NR mice 24 hours before recording. (H-L) Ketamine increased frequency and amplitude of sEPSC in Res but not NR mice. (H) Representative traces of sEPSCs. Scale bar, 50 pA, and 1 s. (I) Frequency. Res(Veh vs Ket), p = 0.0002; NR(Veh vs Ket), p = 0.97, Res+Veh vs NR+Veh, p = 0.84, Res+Ket vs NR+Ket, p = 0.014. F_(3,43)_ = 8.02. (J) Amplitude. Res (Veh vs Ket), p = 0.0009; NR (Veh vs Ket), p > 0.99, Res+Veh vs NR+Veh, p = 0.95, Res+Ket vs NR+Ket, p = 0.0001. F_(3,44)_ = 11.3. (K) Input resistance. Res (Veh vs Ket), p = 0.99; NR (Veh vs Ket), p = 0.53, Res+Veh vs NR+Veh, p = 0.24, Res+Ket vs NR+Ket, p = 0.902. F_(3,52)_ = 1.94. (L) Membrane potential. Res (Veh vs Ket), p = 0.98; NR (Veh vs Ket), p = 0.46, Res+Veh vs NR+Veh, p = 0.99, Res+Ket vs NR+Ket, p = 0.72. F_(3,52)_ = 0.78. n = 11-19 neurons from 3 mice of each group. One-way ANOVA, followed by Tukey’s post hoc. Data, mean ± SEM. *** p < 0.001, * p < 0.05, NS, not significant.

We monitored the calcium activity of same excitatory neurons before and after ketamine treatment (blue cells in Figure 5D). To eliminate potential variability of calcium signals in live animals, we focused on calcium activities during the feeding period in the NSF. As shown in Figure 5E, the frequently and amplitude of calcium transients were increased in Res mice after ketamine, but not in NR mice. Remarkably, in Res mice, normalized calcium activities of PFC neurons were increased by ketamine, compared with Veh controls (z-score of area under the curve normalized with time, 4.0 ± 0.86 with Veh to 4.33 ± 0.97 with ketamine, p < 0.0001, see equation3) (Figure 5F). No difference was detected in calcium activities of PFC neurons between Veh and ketamine groups in NR mice (Veh: 3.95 ± 0.96 vs ketamine: 4.11 ± 0.97, p = 0.26). These results indicate that PFC excitatory neurons in Res mice, but not NR mice, were able to respond to ketamine with increased calcium activities.

Ketamine was known to potentiate the glutamatergic transmission in PFC of human and animals.^14,37^ To investigate cellular mechanisms of NR mice’s inability to respond to ketamine, we next determined whether the ketamine response was altered in NR mice by recording layer 5 PFC pyramidal neurons in whole cell patch configuration. PFC slices were isolated from mice that were treated with ketamine or Veh (Figure 5G). As shown in Figure 5H-5J, both frequency and amplitudes of spontaneous excitatory postsynaptic currents (sEPSC) were higher in the ketamine group, compared with Veh, in Res mice (frequency: 3.6 ± 1.6 Hz with Veh to 10.7 ± 6.1 Hz with ketamine, p = 0.0002; amplitudes: 11.5 ± 4.6 pA with Veh to 19.0 ± 6.6 pA with ketamine, p = 0.0009). In contrast, neither the frequency nor amplitude of sEPSC was altered by ketamine in NR mice (frequency: 4.8 ± 2.4 Hz with Veh to 5.5 ± 3.8 Hz with ketamine, p = 0.97; amplitudes: 10.4 ± 2.8 pA with Veh to 10.3 ± 1.3 pA with ketamine, p > 0.99) (Figure 5I-5J). However, the input resistance and membrane potential were not different between Res and NR mice as well as Veh or ketamine administration (Figure 5K-5L). These results indicate that PFC neurons in NR mice were unable to respond to ketamine with increased glutamatergic transmission. A parsimonious explanation of these results and those from in vivo calcium recordings is that inability of NR mice to respond to ketamine may be due to impaired glutamatergic transmission and thus neuronal activity in the PFC, revealing a mechanism of non-responsiveness to ketamine in NR mice.

### GRP78 for glutamatergic transmission

To determine that GRP78 is critical to glutamatergic transmission, we first investigated whether GRP78 is necessary for ketamine to enhance glutamatergic transmission. Grp78(f/w) Res mice were injected with Cre or GFP viruses in the PFC, to reduce GRP78 expression (Figures 3 and 6A). sEPSCs of layer 5 pyramidal neurons were recorded in PFC slices of injected mice (Figure 6A). As shown in Figure 6B-6D, in Res mice injected with GFP virus, sEPSC frequency and amplitudes were higher in ketamine-treated slices, compared with slices treated with Veh (frequency:2.26 ± 1.78 Hz with Veh; 6.32 ± 2.34 Hz with ketamine, p < 0.001; amplitudes: 9.06 ± 1.49 pA with Veh; 12.2 ± 3.22 pA, with ketamine, p < 0.01). In contrast, no difference was detected in sEPSC frequency or amplitudes between GFP with Veh and Cre virus-injected mice with ketamine (frequency: 3.36 ± 1.81 Hz, p = 0.424; amplitudes: 9.60 ± 1.14 pA, p = 0.83). These results demonstrated that GRP78 is required for PFC neurons to respond to ketamine with increased sEPSCs.

**Figure 6.**
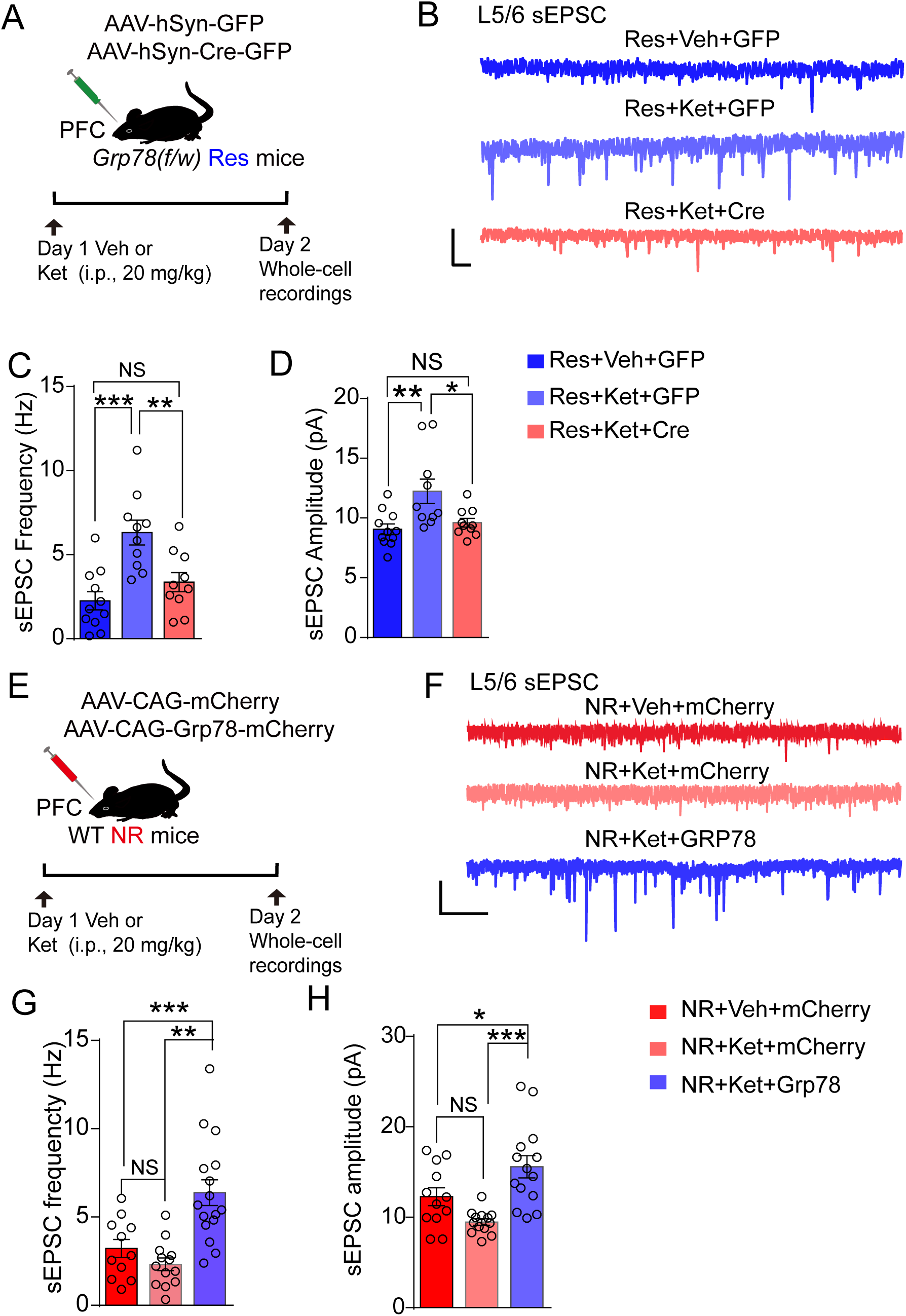
GRP78 for glutamatergic transmission. (A-D) Reducing GRP78 in Res mice blocked the effects of ketamine on sEPSC. (A) Schematic of whole-cell recordings in layer 5 pyramidal neurons of PFC. (B) Representative sEPSC traces from each group. Ketamine was injected 24 hours before recording (20 mg/kg, i.p.). Scale bar, 20 pA, 1 s. (C) Frequency of sEPSC. Res+Veh+GFP vs Res+Ket+GFP, p = 0.0002; Res+Ket+GFP vs Res+Ket+Cre, p = 0.0069, F_(2, 28)_ = 11.5. (D) sEPSC amplitude. Res+GFP+Veh vs Res+Ket+GFP, p = 0.0057; Res+Ket+GFP vs Res+Ket+Cre, p = 0.0269, F_(2, 28)_ = 6.47. n = 12-19 cells from 3 mice for each group. (E-H) Increasing GRP78 in NR mice restored the effects of ketamine on sEPSC. (E) Schematic of experimental design. (F) Representative sEPSC traces from each group. Scale bar, 50 pA, 500 ms. (G) Frequency. NR+Veh+mCherry vs NR+Ket+mCherry, p = 0.58; NR+Ket+mCherry vs NR+Ket+GRP78, p < 0.0001, F_(2, 38)_ = 13.9. (H) Amplitude. NR+Veh+mCherry vs NR+Ket+mCherry, p = 0.17; NR+Ket+mCherry vs NR+Ket+GRP78, p = 0.012, F_(2, 41)_ = 4.72. n = 12-19 cells from 3 mice for each group. One-way ANOVA, followed by Tukey’s *post hoc*. Bars, mean ± SEM. * p < 0.05, ** p < 0.01, ***p < 0.001. NS, not significant.

Next, we tested whether elevating GRP78 in the PFC is sufficient to enable PFC neurons from NR mice to respond to ketamine. NR mice were injected with mCherry or GRP78 viruses, respectively and sEPSCs were recorded in resulting PFC slices (Figure 6E). As shown in Figures 6F-6H, sEPSC frequency and amplitudes in PFC slices of mCherry virus-injected NR mice were similar between ketamine and Veh treatments (frequency: 3.21 ± 1.71 Hz with Veh; 2.32 ± 1.28 Hz with ketamine, p = 0.58; amplitudes: 12.28 ± 3.40 pA with Veh; 9.48 ± 1.28 pA with ketamine, p = 0.17). However, in slices from GRP78 virus-injected mice, ketamine was able to enhance the frequency and amplitudes of sEPSCs, compared mCherry with Veh (frequency: 6.38 ± 2.92 Hz, p < 0.001; amplitudes: 15.6 ± 4.56 pA with GRP78 and ketamine, p = 0.047).

These data showed that increasing GRP78 levels in the PFC of NR mice restored the potentiation of sEPSCs by ketamine. The increase of sEPSCs was associated with raised numbers of spines of PFC neurons that were back filled with biocytin in GRP78 virus-injected NR mice, compared with mCherry virus-injected controls (Figure S8A-S8E). These observations suggest that increased GRP78 expression restored ketamine’s effects in NR mice probably through upregulation of glutamatergic transmission.

### Anti-depressive effects of azoramide by increasing GRP78

GRP78 is a major ER chaperone protein critical for protein quality control of the ER and has been implicated in cell stress such as unfolded protein response.^38,39^ Its level could be increased by azoramide, a hydrophobic small molecule for ER calcium homeostasis, identified by phenotypic screens in response to ER stress (Figure 7A).^43^ We wondered whether azoramide may alter depressive-like behaviors. First, we determined whether it could cross the blood brain barrier (BBB) by injecting mice with azoramide (50 mg/kg, i.p.). Five hours later, mice were sacrificed and perfused extensively with PBS to remove residual blood in the brain (Figure S9A). Cortical tissues were homogenized and subjected to liquid chromatography–mass spectrometry analysis (Figure S9B). As shown in Figure S9C, azoramide was detectable in the cortex of mice injected with azoramide, but not in mice injected with Veh, suggesting that azoramide was able to cross the BBB in agreement with its hydrophobic property. In accord, GRP78 levels were increased in the PFC of azoramide-injected mice, compared with those of mice injected with Veh (Figure 7B-7C).

**Figure 7.**
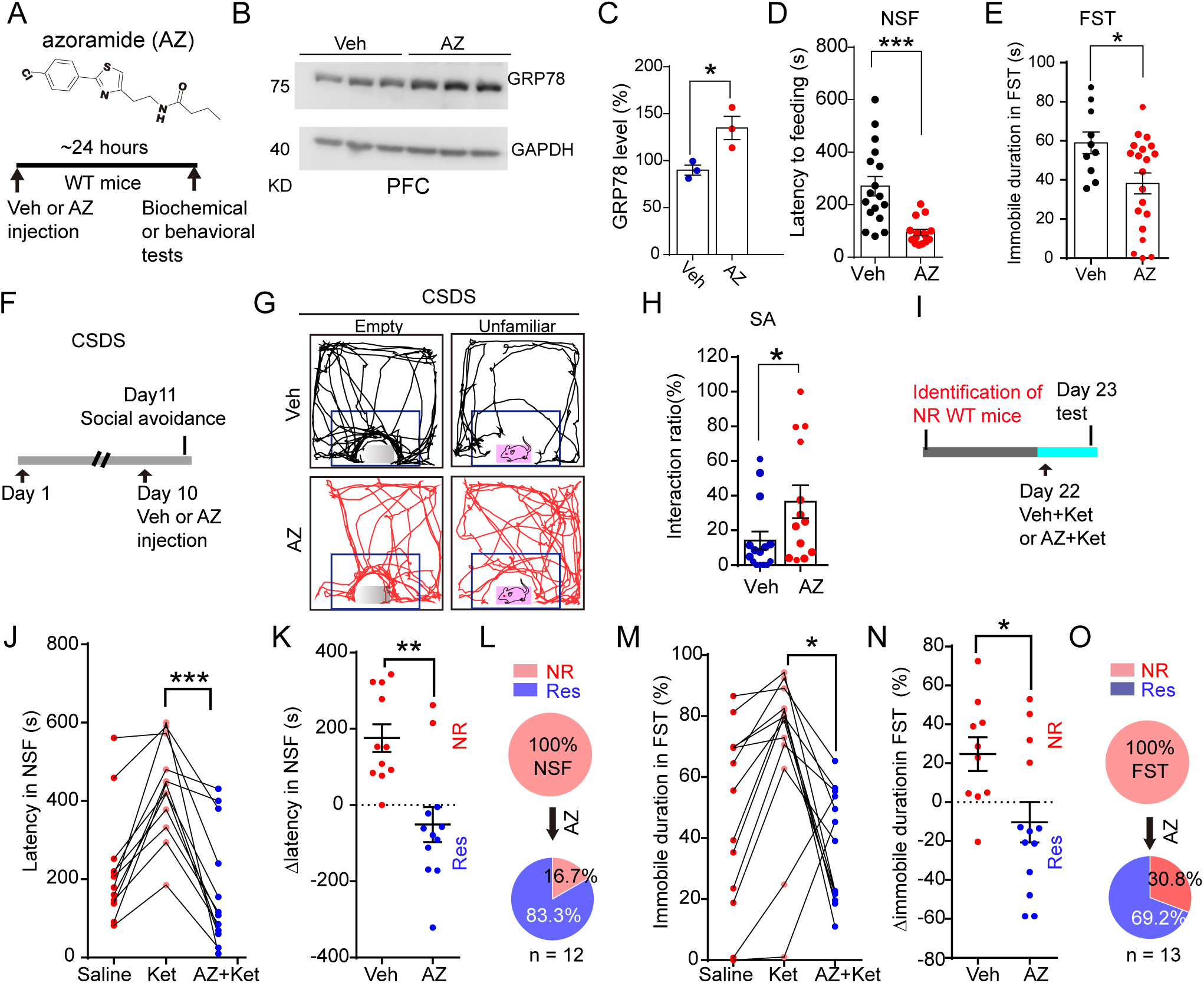
Anti-depressive effects of azoramide by increasing GRP78. (A) Chemical structure of AZ and experimental design. AZ was injected (i.p., 50 mg/kg) 24 hours before experiments. (B-C) Injection of AZ increased the GRP78 level in the PFC. (B) Western blotting of GRP78 in PFC, and GAPDH was used as loading control. (C) Quantification of GRP78 in B. p = 0.029, t = 3.32. n = 3 mice for each group. (D-H) AZ produced antidepression-like effects. (D) Latency in NSF, Veh vs AZ, p < 0.0001. U= 25.0. Mann-Whitney test. (E) Immobile duration in FST. Veh vs AZ, p = 0.021. t = 2.44. t-test. n = 10-20 mice for each group. (F-H) SA after CSDS. (F) Schedule of experiments. (G) Representative traces of animals injected with Veh or AZ in SA. Black, Veh, red, AZ. (H) Interaction ratio in SA. p = 0.042, t = 2.15. T-test. n = 13-15 mice for each group. (I-O) AZ restored NR mice’s response to ketamine in NSF and FST. (I) Experimental design. NR (WT) mice were identified by NSF and FST tests. (J-L) NSF. (J) Latency in NSF in AZ-treated NR mice. Friedman’s F = 20.7, n = 12 mice. (K) Δlatency in NSF in Veh and AZ-treated mice. Veh vs AZ, t = 3.83, p = 0.001, t-test, n = 11-12 mice for each group. (L) Percentage of Res mice from AZ group in Figure 7K (Δlatency < 0). n = 12 mice. Red dots, NR, blue, Res. (M-O) FST. (M) Immobile duration after AZ. Friedman’s F_(12, 24)_ = 1.87, n = 13 mice. Repeated measures ANOVA, followed by Tukey’s *post hoc*. (N) Δimmobile duration after Veh and AZ. T-test, t = 2.49. n = 10-13 mice for each group. (O) Pie chart of NR or Res mice from Figure 7N (AZ group). n = 13 mice. Red dots, NR, blue, Res. * p < 0.05, ** p < 0.01, ***p < 0.001. Data, mean ± SEM.

Next, we subjected azoramide-injected mice to the battery of behavioral tests. Noticeably, azoramide-injected mice showed reduced latency to feeding in NSF, compared with Veh-injected mice (latency to feeding: 270.7 ± 151.9 s with Veh; 94.7 ± 47.2 s with azoramide, p < 0.0001, Figure 7D).

Azoramide also reduced the immobile duration in FST (58.9 ± 17.7% with Veh; 38.2 ± 23.7% with azoramide, p = 0.021, Figure 7E). These results suggest that azoramide may have anti-depressive-like effect in naïve mice. In addition, azoramide increased the interaction ratio of CSDS-stressed mice (14.2 ± 20.1% with Veh; 36.6 ± 34.2% with azoramide, p = 0.042, Figure 7F-7H).

Because azoramide increased GRP78 levels in the PFC (Figure 7B-7C) and because increasing GRP78 levels could enable NR mice to respond to ketamine, we hypothesized that azoramide could also increasing response to ketamine in NR mice. To this end, NR mice were screened out from naïve mice based on response to ketamine, as described in Figure 4G and treated with Veh or azoramide (Figure 7I). As shown in Figure S9D-S9E, NR mice injected with Veh remained nonresponsive to ketamine in NSF and FST, with little changes in latency to feed or immobile duration (latency to feeding: 386.3 ± 129.5 s and 461.7 ± 129.5 s before and after Veh, p > 0.99; immobile duration: 69.8 ± 12.1% and 57.6 ± 22.7% before and after Veh, p = 0.211). In contrast, azoramide-treated NR mice reduced the latency to feeding (430.4 ± 123.9 s and 173.3 ± 151.7 s before and after azoramide, p < 0.0001, Figure 7J), and decreased the immobile duration (70.0 ± 27.2% and 36.9 ± 18.2% before and after azoramide, p = 0.017, Figure 7M). After azoramide treatment, Δlatency in NSF was changed from 175.6 ± 119.7 s with Veh to −51.0 ± 159.4 s with azoramide (p = 0.001), and Δimmobile duration in FST from 24.7 ± 27.5% with Veh to −10.3 ± 37.4% with azoramide (p = 0.022) (Figure 7K and 7N). Overall, 83.3% of NR mice in NSF, and 69.2% in FST became responsive to ketamine (Figure 7L and 7O). These results demonstrate that azoramide could enable NR mice to respond to ketamine likely by increasing GRP78.

We have shown that increasing GRP78 expression enhanced spine numbers and potentiated sEPSC amplitude and frequency in the PFC (Figure 6 and Figure S8). If azoramide alleviates depressive-like behaviors by increasing GRP78, we anticipated that azoimide would increase the spine numbers and sEPSCs. To test this, Thy1-GFP mice were injected with Veh or azoramide and PFC sections were examined for spine morphology (Figure S10A). As shown in Figure S10B-S10D, the spine density of PFC pyramidal neurons was increased in azoramide treated mice, compared with Veh-treated mice. These findings are consistent with our GRP78 gain-of-function studies (Figure S8).

Electrophysiological recordings revealed that azoramide increased the frequency and amplitude of sEPSCs in PFC pyramidal neurons, indicative of potentiation of glutamatergic transmission (Figure S10E-S10H). Together, these results support the model that azoramide exerts antidepressant-like effects through upregulating GRP78, which enhances glutamatergic transmission.

## Discussion

In a non-biased screen for DEGs between mice that respond to ketamine and those that do not, this paper provides evidence that GRP78 plays a critical role in regulating ketamine’s antidepressant effects. First, GRP78 was reduced in mice that were unable to respond to ketamine, ranked the top DEG based on statistical significance (p =1.25E-11). The reduction was confirmed at protein levels by western blot analysis. Second, reducing GRP78 levels in PFC neurons increased latency to feeding in NSF test, immobile duration in FST, and reduced interaction ratio after CSDS, indicating that loss of GRP78 in PFC might induce depression-like behaviors. Third, GRP78 deficiency prevented naïve as well as CSDS-stressed mice from responding to ketamine, suggesting a necessary role of GRP78 in executing ketamine’s anti-depressive like effect. Fourth, remarkably, reducing GRP78 levels in the PFC was able to reduce the responses of ketamine in Res mice whereas enhancing its levels enabled NR mice to respond to ketamine. Finally, treating mice with azoramide, a small chemical that increases GRP78, produced antidepressant-like effects in naïve as well as NR mice. These results demonstrate that altering GRP78 levels not only regulate depressive-like behaviors in naïve mice but also alter the responses of NR and Res mice. A parsimonious explanation of these observations is that GRP78 could be a target of ketamine.

GRP78 is a chaperone protein in the ER that is implicated in managing ER stress and the unfolded protein response (UPR). Our findings that GRP78 regulates depressive-like behaviors and ketamine’s response suggest a role of ER stress and protein folding in regulating depressive-like behaviors and in ketamine’s effects. Indeed, among 1375 down-regulated DEGs, 180 (15.2%) genes were ER proteins, and 28 (2.4%) are involved in ER stress and protein folding. In particular, XBP1 and MANF were both reduced in NR mice. XBP1 is a transcription factor that is induced by ER stress and increases the expression of ER proteins including GRP78 to enhance protein folding and promote ER-associated degradation.^44,45^ MANF is induced during ER stress, likely under the control of XBP1;^46,47^ it does not fold proteins, like GRP78 but might modulate UPR and protect cells from apoptosis.^48,49^ MANF may also be secreted and via cell surface receptors, activates PI3K/Akt and ERK pathways to enhance cell resilience to stress. Dysregulation of XBP1, MANF, and/or GRP78 has been linked to metabolic diseases such as diabetes, hepatic steatosis, and cancer^50–53^ and neurodegenerative disorders including Alzheimer’s and Parkinson’s diseases.^53,54^ ER stress has been increasingly implicated in mental disorders.

ER stress proteins are increased in temporal cortex of depressed subjects who die by suicide and in patients with stress-related mental disorders.^26,27^ On the other hand, the downregulation of XBP1 and GRP78 was reported in monozygotic twins with bipolar disorders.^28^ Moreover, mood stabilizing drugs or antidepressants such as fluoxetine, lithium, valproate could cause ER stress or increase GRP78 in cultured cortical cells.^28,55,56^

In classic cell stress, GRP78 serves both as a sensor and as an effector to clear misfolded proteins.^30,38^ We find that ketamine could enhance the levels of GRP78 in Res mice, but not NR mice, in accord with observations with cultured cells (Figure S5G).^32,57^ This could suggest that NR mice may not be able to sense the effects of ketamine to elevate GRP78. Intriguingly, XBP1 has been shown to bind to the promoter of GRP78, to increase its expression in response to ER stress.^45^ As shown in Figure 1O, Xbp1 is reduced in the PFC of NR mice; therefore, it might be possible that the regulatory mechanisms of Xbp1 expression may be a target of ketamine. On the other hand, the results of loss-of-function and gain-of-function studies described above indicate that GRP78 is not only required for normal mood conditions, but also sufficient to enable NR mice to respond to ketamine. Together, these results could suggest that GRP78 may be a sensor and effector of ketamine. How GRP78 acts to execute ketamine’s antidepressant effects remains unclear. Because ER stress has been detected in MDD patients,^27,58^ elevated GRP78 levels may help alleviate ER stress. Alternatively, GRP78 could protect neurons and regulate neural plasticity by activating PI3K–AKT or mTOR pathways.^30,59,60^ Finally, ketamine’s antidepressant effects are rapid and long-lasting. It is unlikely that ER stress response and GRP78 are involved in the fast effects, considering the time needed for GRP78 expression. Whether GRP78 activation by ketamine is dependent on membrane targets known for its acute effects including NMDAR and potassium channels warrants future studies.

In sum, our work uncovers a previously unrecognized role for ER stress signaling, particularly GRP78, in determining individual responses to ketamine. These findings not only advance our understanding of ketamine’s mechanism of action but also identify GRP78 and ER stress modulation as promising targets for developing novel antidepressants. The ability of GRP78 expression and azoramide to reverse ketamine nonresponse underscores the translational relevance of these pathways as a potential target for therapeutic intervention.

## Materials and Methods

### Mice

Experiments with animals were approved by the Institutional Animal Care and Use Committee of Case Western Reserve University. Mice were housed on a 12 hours controlled light/dark cycle with free access to food and water. WT mice with C57Bl/6j background were used for behavioral assays. In the screening of Res and NR mice, sibling from both genders were used. Grp78^f/f^ mice (Strain #: 035444) were from the Jackson Labs. Genotyping primers of Grp78^f/f^ were: 5’-AGA TGC TGG CAC TAT TGC TG, 5’-ACC CAG GTC AAA CAC AAG GA, that yields a 218 bp product for WT allele and a 260 bp product for Grp78 floxed allele. Genotyping of Nex-Cre mice were performed using the following primers: 5’-GAG TCC TGG AAT CAG TCT TTT TC, 5’-AGA ATG TGG AGT AGG GTGAC, and 5’-AAG CAT AAC CAG TGA AAC AG. WT allele was identified by a 770 bp PCR product, 525 bp for Nex-Cre mutant. Both genders were used for the experiments.

### Behavioral tests

C57Bl/6j background mice 2 to 6 months-old (male and female in some experiments) were used for the behavioral tests. Behavioral tests were performed at 10:00 to 14:00 AM in the light cycle. Health and condition of the mice were examined routinely. Experimenters were blind to the manipulations as well as genotype. Before tests, animals were habituated to the test room for ∼1 hour.^61^

#### Novelty suppressed feeding test

NSF experiments were performed as described before.^62,63^ Mice were food deprived in their home cage for 16 to 24 hours. Food pellet (∼ 3-3.5 cm in length) was fixed in a petri dish (15 cm in diameter) at the center of the open-field arena (50 x 50 cm, 800 lux). In the first NSF test, the floor was covered with corn cob bedding. In the second NSF after ketamine, the floor was not covered with any bedding, and in the NSF3 the floor was covered with alpha cellulose bedding. A video camera was place above the testing arena for recording of the behaviors. ΔLatency was calculated according to the following equation:

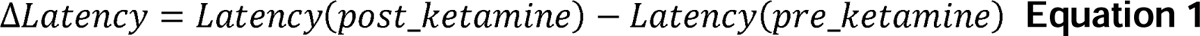

#### CSDS and SA

Male mice were exposed to social defeat stress for 10 days as described before^41^. After CSDS, all the NR and Res mice were infected with Veh. interaction ratio < 100% mice were categorized into susceptible for ketamine treatment 7 days later (Figure S1E). On day 11, the SA was performed in an open field arena (50 × 50 cm) with a mesh cage on one side under darkness. Test will be in two sessions, either empty mesh cage or in the presence of an unfamiliar aggressor in the cage.

#### Forced swimming Test

FST were performed in a tank with a diameter of 20 cm. Water temperature was set to ∼25 ± 1°C, and animals were put in a cage on a heater to dry the fur. Tests were performed for 6 mins, only the last 4 mins was used for analysis. In the first test of FST, tank was not covered; and in the second NSF wall of the tank was covered with white paper (300 lux); the third FST wall was covered with color paper, and illumination was increased to 500 lux. The total immobile duration, and Δfmmobile duration was calculated:

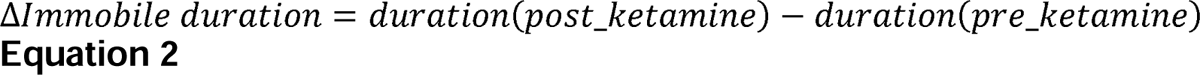

### Tissue collections and western blotting

Brain tissues were collected from coronal sections with the following coordinate: PFC sections (1000 µm, around 2.0 mm to 1.00 mm anterior to bregma).

Sections were cut using a vibratome (VT1000S, Leica Biosystems, USA) in oxygenated PBS buffer on ice. PFC tissues were then dissected under a fluorescent stereo microscope (MVX10, Olympus, Japan). The tissues was homogenized in RIPA buffer [150 mM NaCl, 50 mM Tris-HCl (pH 7.4), 2 mM EDTA, 0.1% SDS, 1 mM phenylmethylsulfonyl fluoride (PMSF), 0.5% sodium deoxycholate, 5 mM NaF, 2 mM NalJVOlJ, phosphorylation inhibitors, and protease inhibitors]. Samples were centrifuged at 12,000 × g for 15 minutes at 4°C to remove debris. Protein concentrations were measured using the Pierce™ BCA Protein Assay Kit (23225, Thermo Fisher Scientific, MA). Samples (17-20 μg) were resolved by SDS-PAGE, transferred to nitrocellulose membranes, and probed with antibodies (see Key Resources Table).

### RNA sequencing and data analysis

Three mice from each group (Res and NR) were euthanized 24 hours post-injection with ketamine (10 mg/kg, i.p.) for PFC tissue collection, as described above in the western blotting protocol. Total RNA was extracted (New England BioLabs), and complementary DNA (cDNA) libraries were constructed. RNA concentration was quantified using a Nanodrop spectrophotometer (Thermo Fisher Scientific). RNA quality was assessed prior to sequencing. RNA sequencing was performed using 50 base pair paired-end reads (Illumina, San Diego, CA). Raw RNA-seq reads were demultiplexed, aligned, and normalized for downstream analysis. Data processing and differential expression analysis were conducted using R packages.^64^ Briefly, DEG analysis was performed using Voom Limma, and threshold of fold change was set to 1.3. The gene enrichment analysis was performed using the GeneOverlap. Related RNA-seq data was deposited to GEO(release later).

### Slice preparation and electrophysiology

Mice were first euthanized and subsequently underwent cardiac perfusion with ice-cold, oxygenated artificial cerebrospinal fluid (ACSF) (95% O_2_ and 5% CO_2_). The brains were rapidly excised and placed in ice-cold, oxygenated ACSF. The ACSF composition included: 110 mM choline chloride, 2.5 mM KCl, 0.5 mM CaCl_2_, 7 mM MgCl_2_, 1.3 mM NaH_2_PO_4_, 25 mM NaHCO_3_, and 20 mM D-glucose. Coronal slices (300 µm) containing the PFC were sectioned using a vibratome (VT1200S, Leica, Germany). The slices were then incubated in oxygenated ACSF for ∼30 minutes at ∼34°C. ACSF used for incubation and recording contained 125 mM NaCl, 2.5 mM KCl, 2 mM CaCl_2_, 1.3 mM MgCl_2_, 1.3 mM NaH_2_PO_4_, 25 mM NaHCO_3_, and 10 mM D-glucose. Following a 60-minute recovery period at room temperature, each slice was transferred to a submerged recording chamber and continuously perfused with oxygenated ACSF at a flow rate of 3.0 ml/min, maintained at approximately 33°C. Slices were visualized using infrared video microscopy with differential interference contrast optics under a microscope (BX51W1, Olympus). L5/6 pyramidal neurons in the PFC were identified based on their distinct location and morphology. Patch electrodes were fabricated from borosilicate glass capillaries (BF150-86-10, Sutter Instruments), with resistances ranging approximately from 3 MΩ, and were filled with the appropriate internal solution. The internal solution for sEPSC recording contained the following: 25 mM cesium methanesulfonate, 5 mM CsCl, 10 mM HEPES, 4 mM MgCl_2_, 4 mM Na_2_ATP, 0.3 mM Na_3_GTP, 10 mM Tris-phosphocreatine and 0.2 mM EGTA, 5 mM QX314. Bicuculline were added to the recording ACSF to block GABAergic transmission.

### AAVs injections

All the AAVs and drugs used for stereotactic injections were listed in Key Resources Table. dTo minimize the toxicity of the AAV solvent, AAVs carrying Cre, GFP, mCherry or GRP78-mCherry used for in vivo experiments were optimized by serial dilution prior to injection. AAV injection surgeries were performed following protocols from previous studies.^61^ Briefly, mice (including WT and Grp78f/w, Nex-Cre) were anesthetized with 2.5% isoflurane in a 30% OlJ: 70% NlJ gas mixture and placed on a heating pad to maintain body temperature at 37°C. The animals were then secured in a stereotactic apparatus for the stereotaxic surgery. Holes targeting the PFC (AP: +1.8 mm, ML: 0.35 mm, DV: 1.5 mm) were drilled, and AAVs were delivered into the target brain region via a fine glass electrode connected to an infusion pump, with an infusion rate of 0.05 µl/min (KDS101, KD Scientific). Afterward, the skin was sutured, and the animals were allowed to recover for at least one week. Subsequently, the mice were subjected to behavioral and electrophysiological experiments.

### Quantitation of Azoramide in mouse plasma and cortex Using HPLC On-line Tandem Mass Spectrometry (LC/MS/MS)

#### Sample Preparation

A 20 µl sample was mixed with 80 µl of methanol containing the internal standard phenylalanine-d5 and then centrifuged at 12,000 rcf for 10 minutes. Following centrifugation, 40 µl of the supernatant was transferred to an HPLC vial for injection. Standard solutions of azoramide were prepared at concentrations of 0, 1, 5, 10, 20, 100, 500, and 2000 ng/ml, each containing phenylalanine-d5 at 1000 ng/ml. The analysis of azoramide compounds was conducted using a triple quadrupole mass spectrometer (Quantiva, Thermo Fisher Scientific, Waltham, MA, USA) coupled to the outlet of an UHPLC system (Vanquish, Thermo Fisher Scientific, Waltham, MA, USA), which included an auto sampler with a refrigerated sample compartment and inline vacuum degasser. A 5 µl volume of the supernatant was injected into a C18 column (Gemini, 2.1 × 150 mm, 3 µm particle size, 110A Phenomenex) for azoramide separation. The mobile phases were A (water containing 0.2% formic acid) and B (acetonitrile containing 0.2% formic acid). The run began with 20% mobile phase B from 0 to 2 minutes. Solvent B was then increased linearly to 100% from 2 to 8 minutes and held at 100% from 8 to 12 minutes. The column was finally re-equilibrated with 20% B for 8 minutes. The HPLC eluent was injected directly into the triple quadrupole mass spectrometer, and the analytes were ionized in positive ESI mode using Selected Reaction Monitoring (SRM). The SRM transitions (m/z) were 486→371 for azoramide and 169→152 for the internal standard phenylalanine-d5. The ion spray voltage was set at 2.5 kV, with sheath gas at 35 Arb and Aux gas at 20 Arb. The ion transfer tube and vaporizer temperatures were set at 350°C and 250°C, respectively. Ultrapure argon (99.99%) was used as a collision gas at a pressure of 2 millitorr for collision-induced dissociation. Data were processed using XCalibur v.4.6 software, which calculated the peak areas for azoramide, the internal standard phenylalanine-d5, and other compounds. Internal standard calibration curves were used to determine the concentration of in brain samples.

### Surgery for calcium imaging

Surgery for calcium imaging was performed as described before with slight modification.^65^ Mice were anesthetized with isoflurane (2.5% isoflurane in a 30% OlJ: 70% NlJ gas mixture) and secured on a stereotaxic apparatus (David Kopf Instruments). Body temperature was maintained at 37°C throughout the procedure using a heating pad.

#### Lens implantation

For lens implantation experiments, mice expressing GCaMP6f in the PFC were placed on a stereotaxic apparatus. A skin incision was made to expose the skull, and connective tissues were carefully removed using a cotton swab. A craniotomy, approximately 1.0 mm in diameter, was performed to target the PFC (AP +2.1 to −1.1 mm, ML 0 to −1.0 mm). Three anchor screws were placed on the skull. A gradient refractive index (GRIN) lens (Inscopix, diameter: 1 mm; length: 4.2 mm; pitch: 0.5; numerical aperture: 0.5) was held by a lens holder (Inscopix) for implantation. To facilitate the penetration of the GRIN lens into the brain, the superficial brain tissue was carefully removed using a 26-gauge blunt tip needle connected to a vacuum pump. Blood was continuously rinsed with sterile PBS to prevent clotting in the hole. After confirming there was no active hemorrhage, the hole was filled with PBS. The GRIN lens, fixed in the lens holder, was inserted through the hole targeting the PFC and gently lowered into the PFC (DV 1.50 mm) at a speed of 0.2 mm/min. After cleaning the skull, the lens was secured to the anchor screws with dental cement. Silicone elastomer (KWIK-CAST, WPI, FL) was applied to cover the exposed end of the lens. Following the surgery, the animal was placed back into its home cage and allowed to recover for 3 weeks.

#### Baseplating

After recovery, mice with implanted lenses underwent a second implantation of base plates for imaging. Since base plating is a non-invasive procedure, the animals were lightly anesthetized (0.5–1% isoflurane) and placed on the stereotaxic apparatus. The miniscope was connected to a holder attached to the manipulator of the stereotaxic apparatus. First, the miniscope was lowered onto the lens, ensuring that no blood clots were present in the field of view (FOV). The desired focus was then achieved by adjusting the height of the miniscope. Finally, the base plate was secured to the rest of the implant using dental cement, and a protective cap was attached to the base plate and secured with screws.

### Calcium imaging data acquisition

#### Collection of data

All calcium imaging data were obtained from defined cell types in the right PFC using a miniaturized integrated fluorescence microscope (nVoke1.0, Inscopix, CA). At the beginning of each imaging session, the miniscope was attached to the baseplate under brief restraint of the animals, without anesthesia. After 45 minutes of habituation in the home cage with the recording setup, animals were transferred to the behavioral apparatus for data collection. Calcium images (970 × 750 pixels) were acquired at a rate of 10.01 fps. To ensure the same field of view (FOV) was imaged across two sessions longitudinally, focus and FOV were not adjusted within the same animal. To minimize photobleaching of the calcium signal from light exposure, images were collected with an LED power of 0.3 mW/mm² and a gain of 1.1. Each imaging session lasted ∼10 minutes, triggered by a TTL signal from a stimulator (Master 9) and synchronized to the behavioral video camera.

### Processing of calcium imaging data

#### Extraction of single cell

The acquired calcium data were preprocessed using Inscopix Data Processing Software (IDPS) First, the imaging data were cropped to the desired field of view (FOV), spatially downsampled, and motion corrected. The preprocessed data were then exported for single-neuron activity extraction using constrained nonnegative matrix factorization for microendoscopic data (CNMF-E)^66^. Single neurons and their fluorescence activity were identified and extracted using the algorithm described previously. For each neuron, the filtered calcium signal trace or deconvolved Ca²⁺events were extracted. To exclude noise and artifacts from blood vessels in the isolated single-cell calcium activity, cells with a size smaller than 20 pixels were rejected for analysis. Each extracted cell was manually inspected for circular spatial contours, and duplicate signal extractions were excluded. Calcium signals corresponding to dendrites or processes, which were more likely to correlate with circular-shaped cell bodies, were also removed from the analysis. Next, the signal-to-noise ratio of the calcium activity was assessed, and Ca²⁺transients were characterized by sharp rises. The analyses were performed using the standard deviation of the filtered fluorescence trace for each cell or the corresponding deconvolved Ca²⁺ transients.

#### Cell registration

Neuron registration from different behavior or stimulation sessions was performed using the CellReg Matlab package as well as visual verification.^67^ Briefly, spatial footprint matrices of cellular activity were obtained from the CNMF-E extraction of single-cell activity. Spatial footprints from all sessions were aligned to a reference coordinate system using rigid-body transformation. The similarities of spatial footprints from neighboring cell pairs across different recording sessions were analyzed by measuring the distances between centroids and spatial correlations. After obtaining the cell registration index for all neurons, the data were manually inspected to exclude false positive and false negative identifications. Subsequently, the calcium activity of paired cells was submitted for further analysis.

### Behavioral analysis in NSF and synchrony with calcium imaging

In the NSF test, behavioral videos were synchronized with calcium imaging data. Behavioral videos were recorded at 40 fps and analyzed as described previously^63^. Briefly, the ImageJ plugin ImageOF (Mouse Phenotype Database; http://www.mouse-phenotype.org/software.html) was used to track the coordinates of the animals’ noses over a 10-minute session^68^. A food pellet was fixed at the center of the open-field arena, and the distance to the food was calculated based on the dynamics of the animal’s coordinates. Then, the animal’s feeding behavior was manually assessed. If the animal’s nose touched the food and it was observed to bite the pellet, this behavior marked the onset of feeding. The period from the beginning of NSF to the onset of feeding was defined as the latency period, and the remaining time was considered the feeding stage. Sniffing, exploration, or passing by the food pellet were not considered non-feeding behaviors.

#### Calcium activity calculation

The calcium activity was analyzed by calculating the calcium activity index. Briefly, ΔF/F was used to compute the area under the curve (AUC) for each neuron’s z-scored calcium activity. The AUC was then normalized by the duration of the specific period. The equation used for the calculation is provided below:

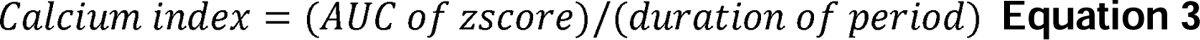

### Statistical analysis

Sample sizes were determined based on power analysis using G*Power 3.1.9.6.^69^ NSF data were analyzed using the Mann Whitney test or Kruskal-Wallis test followed by Dunn’s multiple comparisons test. Same mice before and after treatment was analyzed using Wilcoxon Rank test (NSF, two groups), Friedman test (NSF three groups), paired t-test (FST two groups) and Repeated measures ANOVA (FST three groups). Other analyses were conducted using one-way ANOVA followed by Tukey’s post hoc test for multiple comparisons (3 or more groups) Graphpad Prism 10.0 (Graphpad, CA, USA). Statistical significance was defined as * p < 0.05.

## ACKNOWLEDGMENTS

We thank members of L.M. and W.-C.X. Labs for fruitful discussions.

## Supplementary Figures

**Figure S1.** Limited pre-exposure effects were observed in repeated NSF and FST tests, and NR and Res mice were validated in the CSDS, related to Figure 1. (A) Experimental design of two saline injections tested in two NSF. (B) No alteration of the latency to feeding in NSF. p = 0.584 W = 63.0, n = 36 mice. Wilcoxon matched-pairs rank test. (C-D) Repeated test in FST. (C) Design of experiments. (D) Immobile duration. P = 0.202, t = 1.31 paired t-test. n = 23 mice for each group. (E-G) Susceptible NR and Res mice in CSDS and SA. (E) NR and Res mice were tested with SA before CSDS. Then susceptible NR and Res were identified, and injected with Veh for SA, and 7 days later for Ketamine. Control mice were NR and Res mice without CSDS stress. (F) NR mice in SA. Control, before CSDS. Control vs Veh, 113.2 ± 28.9% vs 37.2 ± 33.9%, p = 0.001. F_(2, 34)_ = 22.5, n = 11-13 for each group. (G) Res mice in SA. Control vs Veh, 126.3 ± 39.2% vs 54.7 ± 16.5%, p < 0.001. F_(2, 29)_ = 13.3, n = 10-11 mice for each group. one-way ANOVA, followed by Tukey’s test. NS, not significant. * p < 0.05, ***p < 0.0001. Data, mean ± SEM.

**Figure S2.** Res and NR mice in NSF and FST with repeated injections of ketamine, related to Figure1. (A-B) Repeated ketamine injections in NSF. (A) Time course of experiments. 10 mg/kg ketamine was injected (i.p.), 24 hours before behavioral tests. (B) Latency. Veh_day 1: NR vs Res, p = 0.76; Ket_day8: NR vs Res, p = 0.004, Ket_day32: NR vs Res, p = 0.029. Interaction, F_(2, 60)_ = 8.75; time, F_(2, 60)_ =2.67; group, F_(1, 30)_ = 4.68. (C-D) FST. (C) Experimental design. (E) Immobile duration in FST. Veh_day 1: NR vs Res, p = 0.86; Ket_day8: NR vs Res, p = 0.042; Ket_day32: NR vs Res, p = 0.0054. Interaction, F_(2, 40)_ = 13.3; time, F_(2, 40)_ =4.38; group, F_(1, 20)_ = 3.71. Two-way ANOVA, followed by Sidak’s post hoc. n = 11-20 mice for each group. * p < 0.05, **p < 0.01. Lines, mean ± SEM.

**Figure S3.** Identification of Res and NR mice by higher doses of ketamine, related to Figure1. (A-C) NSF under higher doses of ketamine. (A) Experimental design. 20 mg/kg ketamine was injected on day 7. (B) Δlatency to feeding in NSF. (C) Pie chart of nonresponders and responders from B. n = 48 mice. (D-F) Nonresponders and responders in FST. (D) Time course of experiments. (E) Distribution of Δimmobile duration in FST. (F) percentage of each population from F. n = 30 mice. Red dots, nonresponders, blue, responders. (G-H) Correlation of Δlatency in NSF and Δimmobile duration in FST. (G) Pearson r correlation analysis, p = 0.008, r = 0.48. Red, NR, blue, Res mice. Dash red line, threshold. (H) Pie chart of NR and Res mice. Red, NR, blue, Res mice. n = 30 mice. Data, mean ± SEM.

**Figure S4.** No difference between Res and NR in baseline as tested in NSF and FST, related to Figure1. (A-B) Baseline behaviors in NSF. (A) Time course of experiments. (B) Latency in NSF. p = 0.22, U = 472, Mann Whitney test. (C-D) FST. (C) Experimental schedule. (D) Immobile duration in FST. p = 0.58, t = 0.56. n = 20-26 mice for each group. T-test. Red dots, NR mice, Blue, Res mice. NS, not significant. Data, mean ± SEM.

**Figure S5.** DEGs and overlapping with ER function, and GRP78 and NMDAR levels in Res and NR mice without ketamine, related to Figure 1. (A) Bar graph showing the total number of DEGs identified in RNA-seq. Light blue, total DEGs, blue, down-regulated, red, up-regulated. (B) Heatmap of DEGs in each sample. (C) Upregulated genes (Top 15, p < 0.05) as analyzed by GO (BP). (D-F) GRP78 and GRIN1 level in Res and NR mice without ketamine. (D) Design of experiment and western blotting. β-Tubulin was used as internal control. (E) Quantification of GRP78. p = 0.16, t = 1.72. n = 3 mice for each group. (F) Quantification of GRIN1. p = 0.61, t = 0.56. T-test. n = 6 mice for each group, bar graph, mean ± SEM. NS, not significant. (G-H) Ketamine increased GRP78 in SH-SY5Y cell line. (G) SH-SY5Y cell line were treated with Veh (saline) or ketamine (5 µm and 10 µm) for 12 hours before harvest. GRP78 and β-Tubulin were analyzed using western blotting. (H) Quantification of GRP78. F_(2, 6)_ = 10.4, Veh vs 5 µM, 100 ± 3.22% vs 168 ± 32.8%, p = 0.017; Veh vs 10 µM, 100 ± 3.22% vs 167 ± 14.9%, p = 0.019. One-way ANOVA followed by Tukey’s multiple comparisons. Data, mean ± SEM. * p < 0.05. NS, not significant.

**Figure S6.** Generation of Grp78 deficiency mice by AAV injection, related to Figure 2 and Figure 3. (A) Gene structure of Grp78(f/w) mice. Exon 5,6, and 7 was flanked by LoxP cassette. After AAV-Cre expression, the Exon 5, 6 and 7 was deleted. (B-C) Reduction of GRP78 in Res mice after AAV injection. (B) Western blotting of Res mice injected with AAV-GFP and Cre in PFC. (C) Quantification of GRP78. p = 0.0031, t = 6.36. T-test. n = 3 mice for each group.

**Figure S7.** Increasing GRP78 in PFC after AAV-GRP78 injection, related to Figure 4. (A) Western blotting of GRP78 in PFC after AAV-mCherry and AAV-GRP78-P2A-mCherry. GRP78-P2A fusion protein was observed in AAV-GRP78 injected mice (upper bands). (B) Quantification of total GRP78. T-test, t = 3.50, p = 0.025. n = 3 mice for each group. * p < 0.05. Data, mean ± SEM.

**Figure S8.** Increasing GRP78 in PFC of NR mice increased the spine density, related to Figure 6. (A) AAV injection. (B) Biocytin labeling of neuronal morphology. Biocytin labeling of PFC pyramidal neurons. Purple, apical, blue, basal spines. (C-E) Increasing of GRP78 in PFC of NR mice increased the spine density in PFC pyramidal neurons. (C) Representative dendritic spines from layer 5 pyramidal neurons of PFC. Scale bar, 10 µm. Upper, basal, lower apical. (D) Basal spine density. p < 0.0001, t = 7.15. (E) Apical. p = 0.0004, t = 3.67. T-test, n = 16-22 cells from 3 mice of each group. One-way ANOVA, followed by Tukey’s post hoc. * p < 0.05, **p < 0.01. Bars, mean ± SEM.

**Figure S9.** Azoramide could penetrate blood-brain barrier and behaviors in Veh-treated NR mice, related to Figure 7. (A-B) Measurement of AZ concentration in cortical tissue after injection. (A) Sampling of cortex from WT mice after AZ injection. Veh or AZ (50 mg/kg) was injected to animals 5 hours before sampling. (B) LC-MS for measurement of AZ. (C) Concentration of AZ in brain cortex. AZ concentration was normalized with protein level. p < 0.0001, t = 116.3. T-test. n = 3 mice for each group. ** p < 0.01. (D) Latency to feeding in NSF after Veh. Saline vs Ket, 286.1 ± 140.5 s vs 386.3 ± 129.5 s, p = 0.0086; Ket vs Veh+Ket, 386.3 ± 129.5 s vs 461.8 ± 142.7, p > 0.99. Friedman test followed by Dunn’s comparison. Friedman F = 15.2. n = 11 mice. (E) Immobile duration in FST after Veh. Saline vs Ket, 32.9 ± 23.3% vs 69.8 ± 12.1%, p = 0.0006; Ket vs Veh+Ket, 69.8 ± 12.1% vs 57.6 ± 22.7%, p = 0.211. Repeated measures ANOVA, followed by Tukey’s comparison. n = 10 mice. NS, not significant, ** p < 0.01. Bar, mean ± SEM.

**Figure S10.** Azoramide injection increased spine density and glutamatergic transmission in PFC, related to Figure 7. (A) Experimental design. Schedule (up panel) and schematic diagram (bottom panel) showing injection of AZ in Thy-1-GFP mice, *i.p.*, AZ, 50 mg/kg. (B) Representative dendritic spine density from Thy1-GFP mice injected with Veh or AZ. Scale bar, 10 µm. (C-D) Spine density in Basal and Apical region. (C) Basal. p = 0.0001, t = 7.17. (D) Apical. p < 0.0001, t = 4.25. n = 51-65 segments of dendrites from 3 mice of each group. T-test. (E-G) AZ increased the sEPSC in PFC pyramidal neurons. (E) Whole-cell recordings of sEPSC in PFC. Pyramidal neurons were identified for recordings. (F) Representative traces of sEPSC from Veh or AZ injected mice. Scale, 20 pA, 0.5 s. Blue, Veh, red, AZ. (G) Frequency. p = 0.0095, t = 2.71. (H) Amplitude. p = 0.04, t = 2.08. n = 22-25 cells from 4 mice of each group. T-test. * p< 0.05, ** p < 0.01, *** p< 0.001. Data, mean ± SEM.

**Table 1.** Downregulated genes in NR mice after ketamine.

**Table 2.** Upregulated genes in NR mice after ketamine.

## References

1. Kessler, R.C., Berglund, P., Demler, O., Jin, R., Koretz, D., Merikangas, K.R., Rush, A.J., Walters, E.E., and Wang, P.S. (2003). The epidemiology of major depressive disorder - Results from the National Comorbidity Survey Replication (NCS-R). Jama-J Am Med Assoc 289, 3095–3105. DOI 10.1001/jama.289.23.3095.

2. Cipriani, A., Furukawa, T.A., Salanti, G., Chaimani, A., Atkinson, L.Z., Ogawa, Y., Leucht, S., Ruhe, H.G., Turner, E.H., Higgins, J.P.T., et al. (2018). Comparative Efficacy and Acceptability of 21 Antidepressant Drugs for the Acute Treatment of Adults With Major Depressive Disorder: A Systematic Review and Network Meta-Analysis. Focus (Am Psychiatr Publ) 16, 420–429. 10.1176/appi.focus.16407.

3. Moncrieff, J., Cooper, R.E., Stockmann, T., Amendola, S., Hengartner, M.P., and Horowitz, M.A. (2023). The serotonin theory of depression: a systematic umbrella review of the evidence. Mol Psychiatry 28, 3243–3256. 10.1038/s41380-022-01661-0.

4. Simon, G.E., Moise, N., and Mohr, D.C. (2024). Management of Depression in Adults: A Review. JAMA 332, 141–152. 10.1001/jama.2024.5756.

5. Gelenberg, A.J., Freeman, M., Markowitz, J., Rosenbaum, J., Thase, M., Trivedi, M., and Van Rhoads, R. (2010). American Psychiatric Association practice guidelines for the treatment of patients with major depressive disorder. Am J Psychiatry 167, 9–118.

6. Hirschfeld, R.M.A. (1999). Efficacy of SSRIs and newer antidepressants in severe depression: Comparison with TCAs. J Clin Psychiat 60, 326–335. DOI 10.4088/JCP.v60n0511.

7. Thomas, L., Kessler, D., Campbell, J., Morrison, J., Peters, T.J., Williams, C., Lewis, G., and Wiles, N. (2013). Prevalence of treatment-resistant depression in primary care: cross-sectional data. Br J Gen Pract 63, e852–858. 10.3399/bjgp13X675430.

8. Entsuah, A.R., Huang, H., and Thase, M.E. (2001). Response and remission rates in different subpopulations with major depressive disorder administered venlafaxine, selective serotonin reuptake inhibitors, or placebo. J Clin Psychiatry 62, 869–877. 10.4088/jcp.v62n1106.

9. Little, A. (2009). Treatment-resistant depression. Am Fam Physician 80, 167–172.

10. Berman, R.M., Cappiello, A., Anand, A., Oren, D.A., Heninger, G.R., Charney, D.S., and Krystal, J.H. (2000). Antidepressant effects of ketamine in depressed patients. Biol Psychiatry 47, 351–354. 10.1016/s0006-3223(99)00230-9.

11. Daly, E.J., Trivedi, M.H., Janik, A., Li, H., Zhang, Y., Li, X., Lane, R., Lim, P., Duca, A.R., Hough, D., et al. (2019). Efficacy of Esketamine Nasal Spray Plus Oral Antidepressant Treatment for Relapse Prevention in Patients With Treatment-Resistant Depression: A Randomized Clinical Trial. JAMA Psychiatry 76, 893–903. 10.1001/jamapsychiatry.2019.1189.

12. Warden, M.R., Selimbeyoglu, A., Mirzabekov, J.J., Lo, M., Thompson, K.R., Kim, S.Y., Adhikari, A., Tye, K.M., Frank, L.M., and Deisseroth, K. (2012). A prefrontal cortex-brainstem neuronal projection that controls response to behavioural challenge. Nature 492, 428–432. 10.1038/nature11617.

13. Yang, Y., Cui, Y., Sang, K., Dong, Y., Ni, Z., Ma, S., and Hu, H. (2018). Ketamine blocks bursting in the lateral habenula to rapidly relieve depression. Nature 554, 317–322. 10.1038/nature25509.

14. Li, N., Lee, B., Liu, R.J., Banasr, M., Dwyer, J.M., Iwata, M., Li, X.Y., Aghajanian, G., and Duman, R.S. (2010). mTOR-dependent synapse formation underlies the rapid antidepressant effects of NMDA antagonists. Science 329, 959–964. 10.1126/science.1190287.

15. Gerhard, D.M., Pothula, S., Liu, R.J., Wu, M., Li, X.Y., Girgenti, M.J., Taylor, S.R., Duman, C.H., Delpire, E., Picciotto, M., et al. (2020). GABA interneurons are the cellular trigger for ketamine’s rapid antidepressant actions. J Clin Invest 130, 1336–1349. 10.1172/JCI130808.

16. Autry, A.E., Adachi, M., Nosyreva, E., Na, E.S., Los, M.F., Cheng, P.F., Kavalali, E.T., and Monteggia, L.M. (2011). NMDA receptor blockade at rest triggers rapid behavioural antidepressant responses. Nature 475, 91–95. 10.1038/nature10130.

17. Lepack, A.E., Fuchikami, M., Dwyer, J.M., Banasr, M., and Duman, R.S. (2014). BDNF Release Is Required for the Behavioral Actions of Ketamine. Int J Neuropsychoph 18. 10.1093/ijnp/pyu033.

18. Pothula, S., Kato, T., Liu, R.J., Wu, M., Gerhard, D., Shinohara, R., Sliby, A.N., Chowdhury, G.M.I., Behar, K.L., Sanacora, G., et al. (2021). Cell-type specific modulation of NMDA receptors triggers antidepressant actions. Mol Psychiatry 26, 5097–5111. 10.1038/s41380-020-0796-3.

19. Lopez, J.P., Lucken, M.D., Brivio, E., Karamihalev, S., Kos, A., De Donno, C., Benjamin, A., Yang, H., Dick, A.L.W., Stoffel, R., et al. (2022). Ketamine exerts its sustained antidepressant effects via cell-type-specific regulation of Kcnq2. Neuron 110, 2283–2298 e2289. 10.1016/j.neuron.2022.05.001.

20. Zanos, P., Moaddel, R., Morris, P.J., Georgiou, P., Fischell, J., Elmer, G.I., Alkondon, M., Yuan, P., Pribut, H.J., Singh, N.S., et al. (2016). NMDAR inhibition-independent antidepressant actions of ketamine metabolites. Nature 533, 481–486. 10.1038/nature17998.

21. Zarate, C.A., Jr., Singh, J.B., Carlson, P.J., Brutsche, N.E., Ameli, R., Luckenbaugh, D.A., Charney, D.S., and Manji, H.K. (2006). A randomized trial of an N-methyl-D-aspartate antagonist in treatment-resistant major depression. Arch Gen Psychiatry 63, 856–864. 10.1001/archpsyc.63.8.856.

22. Szymkowicz, S.M., Finnegan, N., and Dale, R.M. (2014). Failed response to repeat intravenous ketamine infusions in geriatric patients with major depressive disorder. J Clin Psychopharmacol 34, 285–286. 10.1097/JCP.0000000000000090.

23. Bagot, R.C., Cates, H.M., Purushothaman, I., Vialou, V., Heller, E.A., Yieh, L., LaBonte, B., Pena, C.J., Shen, L., Wittenberg, G.M., and Nestler, E.J. (2017). Ketamine and Imipramine Reverse Transcriptional Signatures of Susceptibility and Induce Resilience-Specific Gene Expression Profiles. Biol Psychiatry 81, 285–295. 10.1016/j.biopsych.2016.06.012.

24. Schwarz, D.S., and Blower, M.D. (2016). The endoplasmic reticulum: structure, function and response to cellular signaling. Cell Mol Life Sci 73, 79–94. 10.1007/s00018-015-2052-6.

25. Araki, K., and Nagata, K. (2011). Protein folding and quality control in the ER. Cold Spring Harb Perspect Biol 3, a007526. 10.1101/cshperspect.a007526.

26. Nevell, L., Zhang, K.Z., Aiello, A.E., Koenen, K., Galea, S., Soliven, R., Zhang, C., Wildman, D.E., and Uddin, M. (2014). Elevated systemic expression of ER stress related genes is associated with stress-related mental disorders in the Detroit Neighborhood Health Study. Psychoneuroendocrino 43, 62–70. 10.1016/j.psyneuen.2014.01.013.

27. Bown, C., Wang, J.F., MacQueen, G., and Young, L.T. (2000). Increased temporal cortex ER stress proteins in depressed subjects who died by suicide. Neuropsychopharmacology 22, 327–332. 10.1016/S0893-133X(99)00091-3.

28. Kakiuchi, C., Iwamoto, K., Ishiwata, M., Bundo, M., Kasahara, T., Kusumi, I., Tsujita, T., Okazaki, Y., Nanko, S., Kunugi, H., et al. (2003). Impaired feedback regulation of XBP1 as a genetic risk factor for bipolar disorder. Nat Genet 35, 171–175. 10.1038/ng1235.

29. Grunebaum, M.F., Galfalvy, H.C., Huang, Y.Y., Cooper, T.B., Burke, A.K., Agnello, M., Oquendo, M.A., and Mann, J.J. (2009). Association of X-box binding protein 1 (XBP1) genotype with morning cortisol and 1-year clinical course after a major depressive episode. Int J Neuropsychopharmacol 12, 281–283. 10.1017/S1461145708009863.

30. Lee, A.S. (2014). Glucose-regulated proteins in cancer: molecular mechanisms and therapeutic potential. Nat Rev Cancer 14, 263–276. 10.1038/nrc3701.

31. Rigg, N., Abu-Hijleh, F.A., Patel, V., and Mishra, R.K. (2022). Ketamine-induced neurotoxicity is mediated through endoplasmic reticulum stress in vitro in STHdh cells. Neurotoxicology 91, 321–328. 10.1016/j.neuro.2022.06.004.

32. Cui, L.J., Jiang, X.Y., Zhang, C.J., Li, D.X., Yu, S.Q., Wan, F.C., Ma, Y., Guo, W., and Shan, Z.F. (2019). Ketamine induces endoplasmic reticulum stress in rats and SV-HUC-1 human uroepithelial cells by activating NLRP3/TXNIP aix. Bioscience Rep 39. Artn Bsr20190595 10.1042/Bsr20190595.

33. Santarelli, L., Saxe, M., Gross, C., Surget, A., Battaglia, F., Dulawa, S., Weisstaub, N., Lee, J., Duman, R., Arancio, O., et al. (2003). Requirement of hippocampal neurogenesis for the behavioral effects of antidepressants. Science 301, 805–809. 10.1126/science.1083328.

34. Ma, S., Chen, M., Jiang, Y., Xiang, X., Wang, S., Wu, Z., Li, S., Cui, Y., Wang, J., Zhu, Y., et al. (2023). Sustained antidepressant effect of ketamine through NMDAR trapping in the LHb. Nature 622, 802–809. 10.1038/s41586-023-06624-1.

35. Cryan, J.F., Valentino, R.J., and Lucki, I. (2005). Assessing substrates underlying the behavioral effects of antidepressants using the modified rat forced swimming test. Neurosci Biobehav Rev 29, 547–569. 10.1016/j.neubiorev.2005.03.008.

36. Abdallah, C.G., De Feyter, H.M., Averill, L.A., Jiang, L., Averill, C.L., Chowdhury, G.M.I., Purohit, P., de Graaf, R.A., Esterlis, I., Juchem, C., et al. (2018). The effects of ketamine on prefrontal glutamate neurotransmission in healthy and depressed subjects. Neuropsychopharmacology 43, 2154–2160. 10.1038/s41386-018-0136-3.

37. Moghaddam, B., Adams, B., Verma, A., and Daly, D. (1997). Activation of glutamatergic neurotransmission by ketamine: a novel step in the pathway from NMDA receptor blockade to dopaminergic and cognitive disruptions associatedwith the prefrontal cortex. J Neurosci 17, 2921–2927. 10.1523/JNEUROSCI.17-08-02921.1997.

38. Lee, A.S. (2005). The ER chaperone and signaling regulator GRP78/BiP as a monitor of endoplasmic reticulum stress. Methods 35, 373–381. 10.1016/j.ymeth.2004.10.010.

39. Li, J., Ni, M., Lee, B., Barron, E., Hinton, D.R., and Lee, A.S. (2008). The unfolded protein response regulator GRP78/BiP is required for endoplasmic reticulum integrity and stress-induced autophagy in mammalian cells. Cell Death and Differentiation 15, 1460–1471. 10.1038/cdd.2008.81.

40. Luo, S.Z., Mao, C.H., Lee, B., and Lee, A.S. (2006). GRP78/BiP is required for cell proliferation and protecting the inner cell mass from apoptosis during early mouse embryonic development. Mol Cell Biol 26, 5688–5697. 10.1128/Mcb.00779-06.

41. Berton, O., McClung, C.A., Dileone, R.J., Krishnan, V., Renthal, W., Russo, S.J., Graham, D., Tsankova, N.M., Bolanos, C.A., Rios, M., et al. (2006). Essential role of BDNF in the mesolimbic dopamine pathway in social defeat stress. Science 311, 864–868. 10.1126/science.1120972.

42. Moda-Sava, R.N., Murdock, M.H., Parekh, P.K., Fetcho, R.N., Huang, B.S., Huynh, T.N., Witztum, J., Shaver, D.C., Rosenthal, D.L., Alway, E.J., et al. (2019). Sustained rescue of prefrontal circuit dysfunction by antidepressant-induced spine formation. Science 364. 10.1126/science.aat8078.

43. Fu, S.N., Yalcin, A., Lee, G.Y., Li, P., Fan, J., Arruda, A.P., Pers, B.M., Yilmaz, M., Eguchi, K., and Hotamisligil, G.S. (2015). Phenotypic assays identify azoramide as a small-molecule modulator of the unfolded protein response with antidiabetic activity. Science Translational Medicine 7. ARTN 292ra98 10.1126/scitranslmed.aaa9134.

44. Shuda, M., Kondoh, N., Imazeki, N., Tanaka, K., Okada, T., Mori, K., Hada, A., Arai, M., Wakatsuki, T., Matsubara, O., et al. (2003). Activation of the ATF6, XBP1 and grp78 genes in human hepatocellular carcinoma: a possible involvement of the ER stress pathway in hepatocarcinogenesis. J Hepatol 38, 605–614. 10.1016/s0168-8278(03)00029-1.

45. Yoshida, H., Matsui, T., Yamamoto, A., Okada, T., and Mori, K. (2001). XBP1 mRNA is induced by ATF6 and spliced by IRE1 in response to ER stress to produce a highly active transcription factor. Cell 107, 881–891. 10.1016/s0092-8674(01)00611-0.

46. Lee, A.H., Iwakoshi, N.N., and Glimcher, L.H. (2003). XBP-1 regulates a subset of endoplasmic reticulum resident chaperone genes in the unfolded protein response. Mol Cell Biol 23, 7448–7459. 10.1128/MCB.23.21.7448-7459.2003.

47. Shaffer, A.L., Shapiro-Shelef, M., Iwakoshi, N.N., Lee, A.H., Qian, S.B., Zhao, H., Yu, X., Yang, L., Tan, B.K., Rosenwald, A., et al. (2004). XBP1, downstream of Blimp-1, expands the secretory apparatus and other organelles, and increases protein synthesis in plasma cell differentiation. Immunity 21, 81–93. 10.1016/j.immuni.2004.06.010.

48. Glembotski, C.C., Thuerauf, D.J., Huang, C., Vekich, J.A., Gottlieb, R.A., and Doroudgar, S. (2012). Mesencephalic astrocyte-derived neurotrophic factor protects the heart from ischemic damage and is selectively secreted upon sarco/endoplasmic reticulum calcium depletion. J Biol Chem 287, 25893–25904. 10.1074/jbc.M112.356345.

49. Hakonen, E., Chandra, V., Fogarty, C.L., Yu, N.Y., Ustinov, J., Katayama, S., Galli, E., Danilova, T., Lindholm, P., Vartiainen, A., et al. (2018). MANF protects human pancreatic beta cells against stress-induced cell death. Diabetologia 61, 2202–2214. 10.1007/s00125-018-4687-y.

50. Petrova, P., Raibekas, A., Pevsner, J., Vigo, N., Anafi, M., Moore, M.K., Peaire, A.E., Shridhar, V., Smith, D.I., Kelly, J., et al. (2003). MANF: a new mesencephalic, astrocyte-derived neurotrophic factor with selectivity for dopaminergic neurons. J Mol Neurosci 20, 173–188. 10.1385/jmn:20:2:173.

51. Lee, A.H., Heidtman, K., Hotamisligil, G.S., and Glimcher, L.H. (2011). Dual and opposing roles of the unfolded protein response regulated by IRE1alpha and XBP1 in proinsulin processing and insulin secretion. Proc Natl Acad Sci U S A 108, 8885–8890. 10.1073/pnas.1105564108.

52. Lee, A.S. (2007). GRP78 induction in cancer: therapeutic and prognostic implications. Cancer Res 67, 3496–3499. 10.1158/0008-5472.CAN-07-0325.

53. Lebeaupin, C., Vallee, D., Hazari, Y., Hetz, C., Chevet, E., and Bailly-Maitre, B. (2018). Endoplasmic reticulum stress signalling and the pathogenesis of non-alcoholic fatty liver disease. J Hepatol 69, 927–947. 10.1016/j.jhep.2018.06.008.

54. Gerakis, Y., and Hetz, C. (2018). Emerging roles of ER stress in the etiology and pathogenesis of Alzheimer’s disease. FEBS J 285, 995–1011. 10.1111/febs.14332.

55. Wang, J.-F., and Young, L.T. (2004). Regulation of molecular chaperone GRP78 by mood stabilizing drugs. Clinical Neuroscience Research 4, 281–288. 10.1016/j.cnr.2004.09.007.

56. Shao, S., Zhuang, X., Zhang, L., and Qiao, T. (2022). Antidepressants Fluoxetine Mediates Endoplasmic Reticulum Stress and Autophagy of Non-Small Cell Lung Cancer Cells Through the ATF4-AKT-mTOR Signaling Pathway. Front Pharmacol 13, 904701. 10.3389/fphar.2022.904701.

57. Józwiak-Bebenista, M., Sokolowska, P., Siatkowska, M., Panek, C.A., Komorowski, P., Kowalczyk, E., and Wiktorowska-Owczarek, A. (2022). The Importance of Endoplasmic Reticulum Stress as a Novel Antidepressant Drug Target and Its Potential Impact on CNS Disorders. Pharmaceutics 14. ARTN 846 10.3390/pharmaceutics14040846.

58. Ii Timberlake, M., and Dwivedi, Y. (2019). Linking unfolded protein response to inflammation and depression: potential pathologic and therapeutic implications. Mol Psychiatry 24, 987–994. 10.1038/s41380-018-0241-z.

59. Fu, Y., Wey, S., Wang, M., Ye, R., Liao, C.P., Roy-Burman, P., and Lee, A.S. (2008). Pten null prostate tumorigenesis and AKT activation are blocked by targeted knockout of ER chaperone GRP78/BiP in prostate epithelium. Proc Natl Acad Sci U S A 105, 19444–19449. 10.1073/pnas.0807691105.

60. Wey, S., Luo, B., Tseng, C.C., Ni, M., Zhou, H., Fu, Y., Bhojwani, D., Carroll, W.L., and Lee, A.S. (2012). Inducible knockout of GRP78/BiP in the hematopoietic system suppresses Pten-null leukemogenesis and AKT oncogenic signaling. Blood 119, 817–825. 10.1182/blood-2011-06-357384.

61. Cui, W., Gao, N., Dong, Z., Shen, C., Zhang, H., Luo, B., Chen, P., Comoletti, D., Jing, H., Wang, H., et al. (2021). In trans neuregulin3-Caspr3 interaction controls DA axonal bassoon cluster development. Curr Biol 31, 3330–3342 e3337. 10.1016/j.cub.2021.05.045.

62. Samuels, B.A., and Hen, R. (2011). Novelty-Suppressed Feeding in the Mouse. In Mood and Anxiety Related Phenotypes in Mice: Characterization Using Behavioral Tests, Volume II, T.D. Gould, ed. (Humana Press), pp. 107–121. 10.1007/978-1-61779-313-4_7.

63. Cui, W., Mizukami, H., Yanagisawa, M., Aida, T., Nomura, M., Isomura, Y., Takayanagi, R., Ozawa, K., Tanaka, K., and Aizawa, H. (2014). Glial dysfunction in the mouse habenula causes depressive-like behaviors and sleep disturbance. J Neurosci 34, 16273–16285. 10.1523/JNEUROSCI.1465-14.2014.

64. Gentleman, R.C., Carey, V.J., Bates, D.M., Bolstad, B., Dettling, M., Dudoit, S., Ellis, B., Gautier, L., Ge, Y., Gentry, J., et al. (2004). Bioconductor: open software development for computational biology and bioinformatics. Genome Biol 5, R80. 10.1186/gb-2004-5-10-r80.

65. Lui, J.H., Nguyen, N.D., Grutzner, S.M., Darmanis, S., Peixoto, D., Wagner, M.J., Allen, W.E., Kebschull, J.M., Richman, E.B., Ren, J., et al. (2021). Differential encoding in prefrontal cortex projection neuron classes across cognitive tasks. Cell 184, 489–506 e426. 10.1016/j.cell.2020.11.046.

66. Zhou, P., Resendez, S.L., Rodriguez-Romaguera, J., Jimenez, J.C., Neufeld, S.Q., Giovannucci, A., Friedrich, J., Pnevmatikakis, E.A., Stuber, G.D., Hen, R., et al. (2018). Efficient and accurate extraction of in vivo calcium signals from microendoscopic video data. Elife 7. 10.7554/eLife.28728.

67. Sheintuch, L., Rubin, A., Brande-Eilat, N., Geva, N., Sadeh, N., Pinchasof, O., and Ziv, Y. (2017). Tracking the Same Neurons across Multiple Days in Ca(2+) Imaging Data. Cell Rep 21, 1102–1115. 10.1016/j.celrep.2017.10.013.

68. Cui, W., Aida, T., Ito, H., Kobayashi, K., Wada, Y., Kato, S., Nakano, T., Zhu, M., Isa, K., Kobayashi, K., et al. (2020). Dopaminergic Signaling in the Nucleus Accumbens Modulates Stress-Coping Strategies during Inescapable Stress. J Neurosci 40, 7241–7254. 10.1523/JNEUROSCI.0444-20.2020.

69. Faul, F., Erdfelder, E., Lang, A.G., and Buchner, A. (2007). G*Power 3: a flexible statistical power analysis program for the social, behavioral, and biomedical sciences. Behav Res Methods 39, 175–191. 10.3758/bf03193146.

