## Supplementary figures and images for "Endoplasmic reticulum stress protein GRP78 for ketamine’s antidepressant effects"

### Supplemental Figures1-10

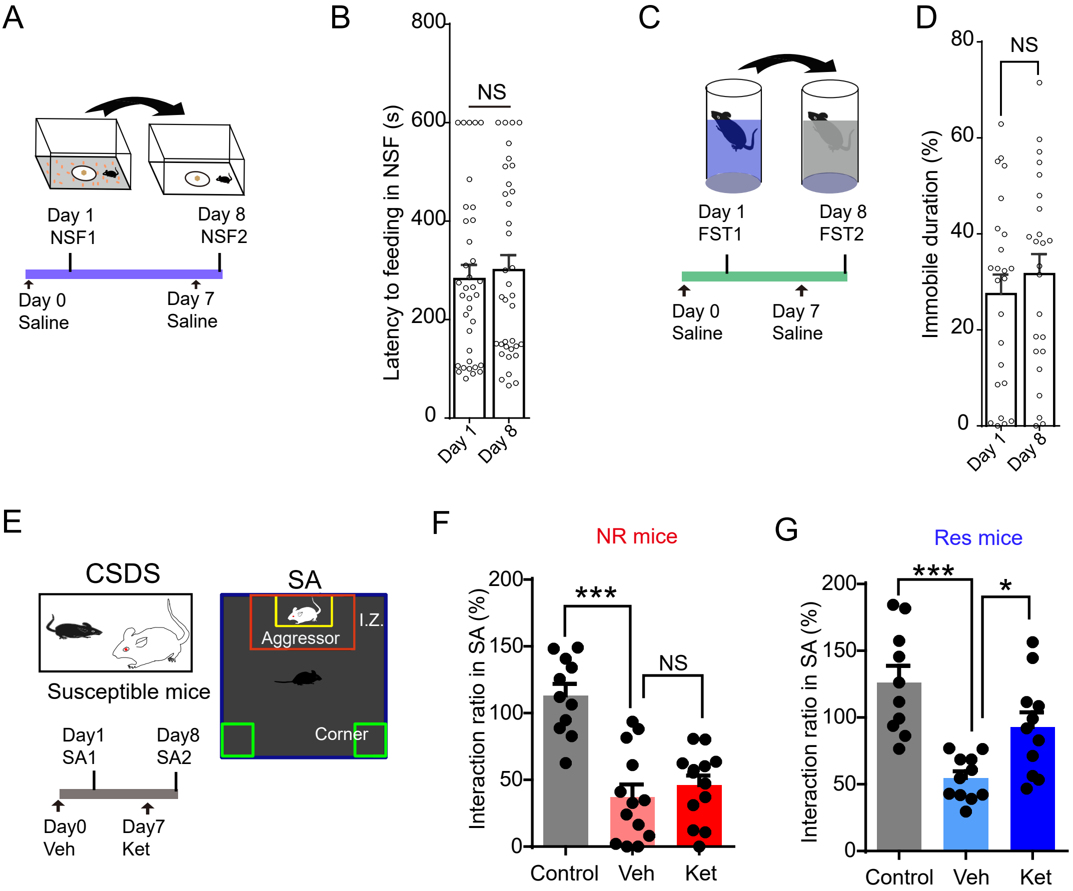

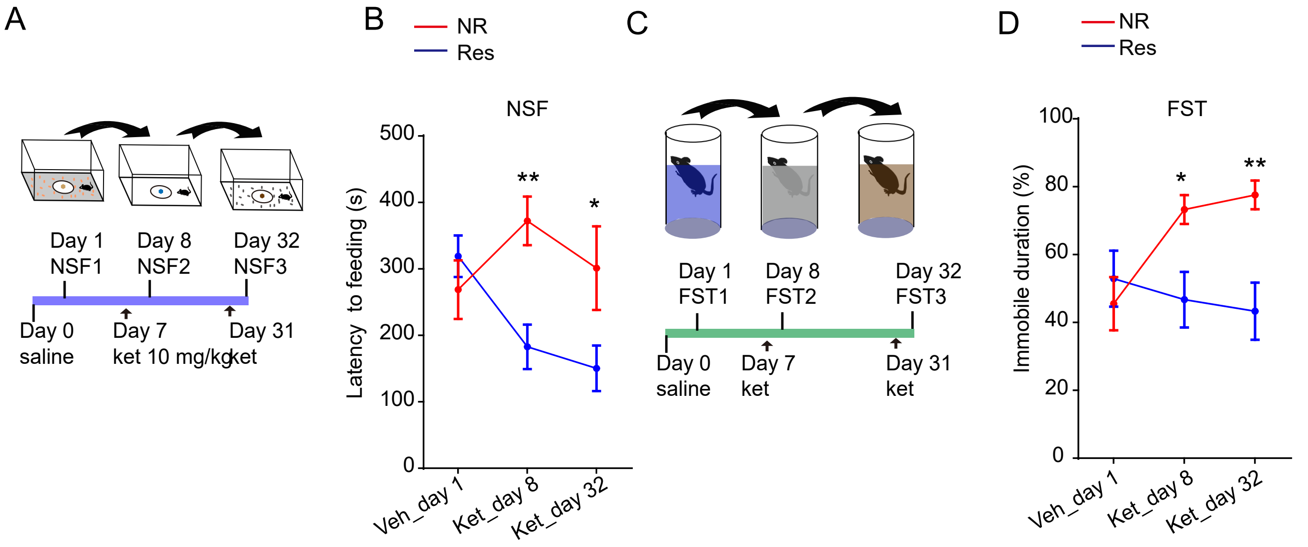

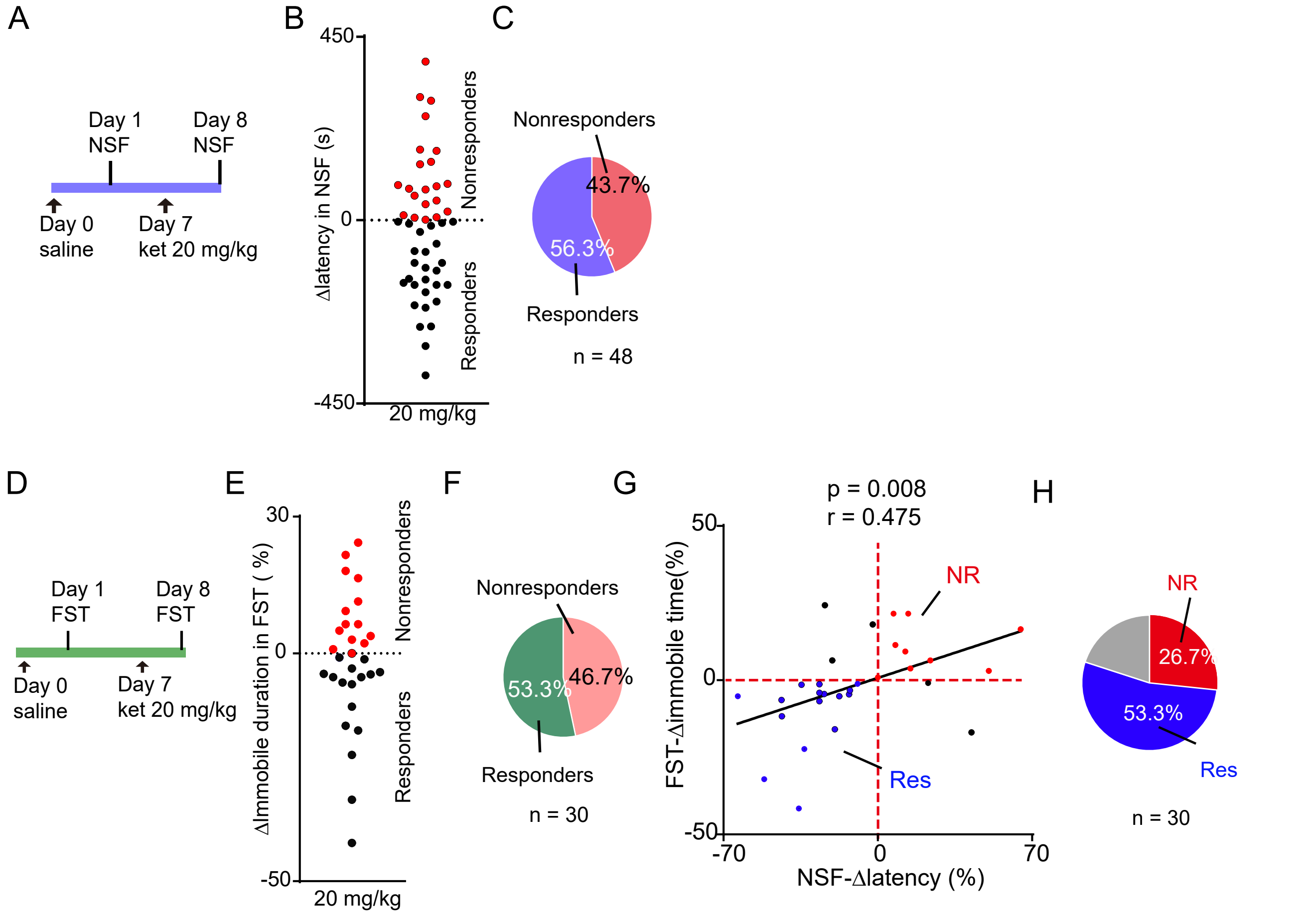

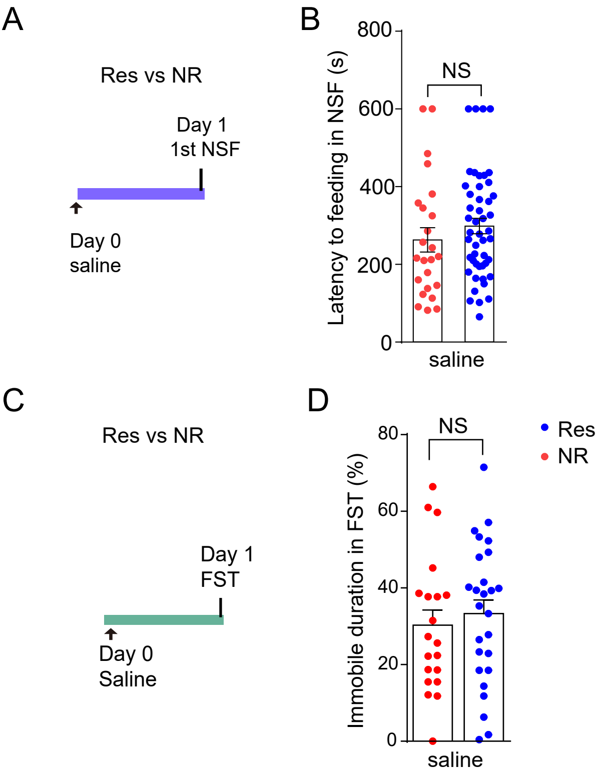

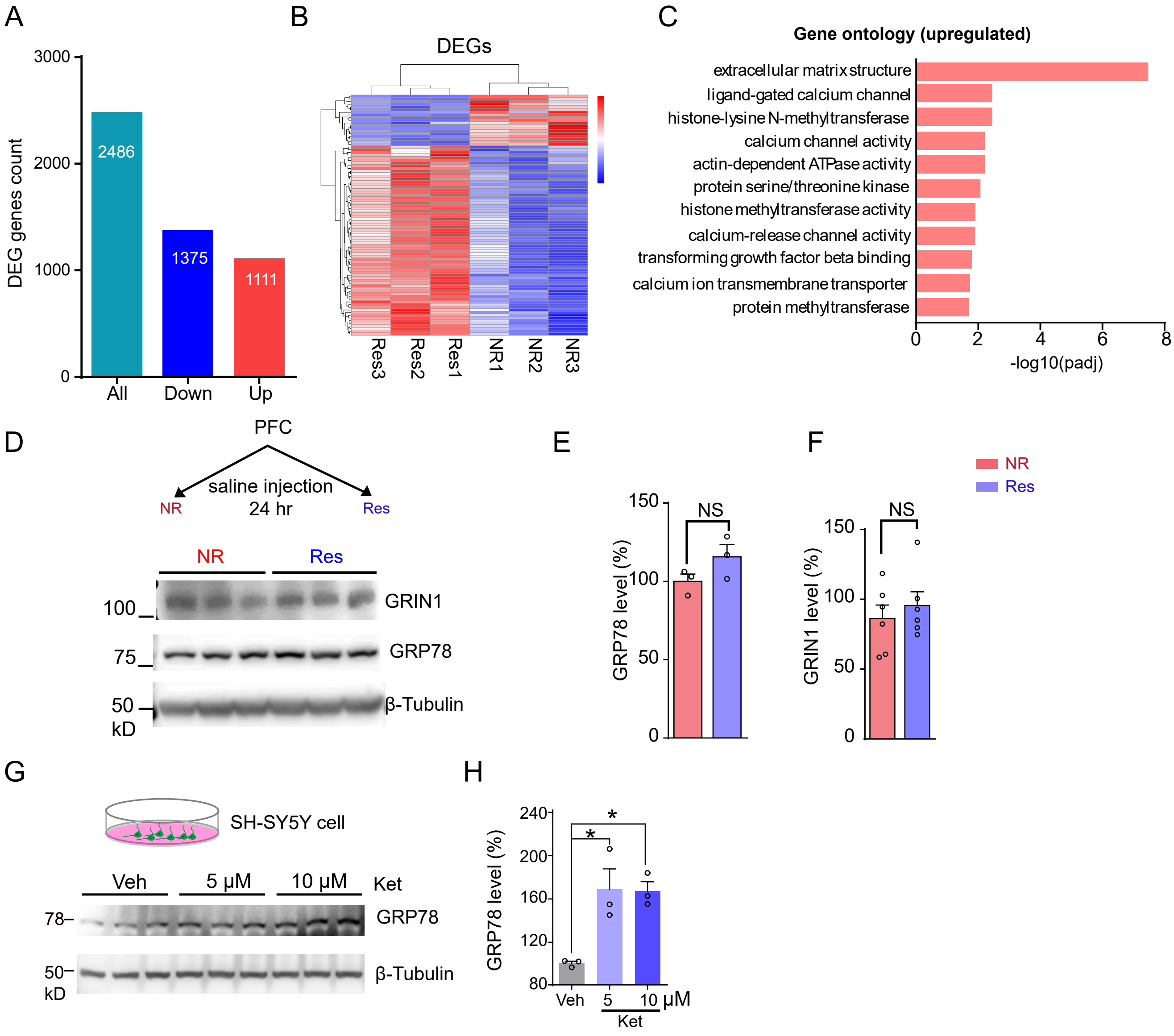

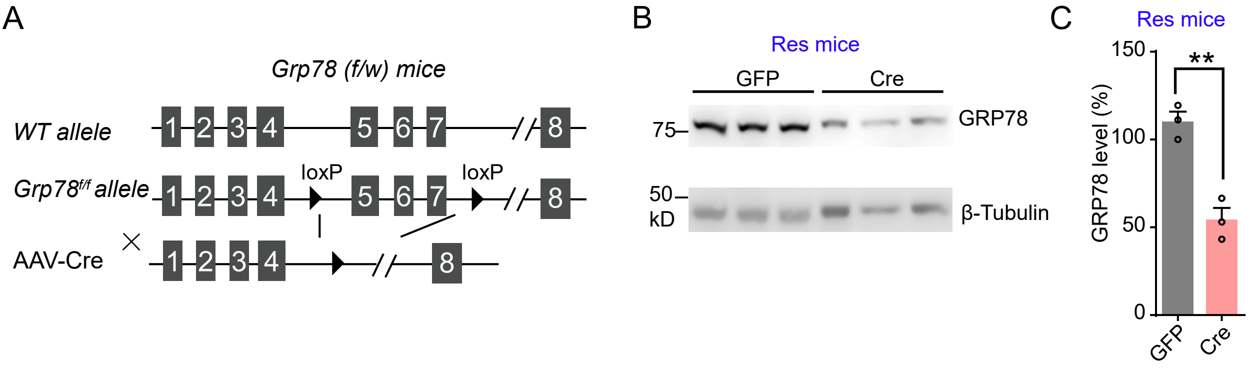

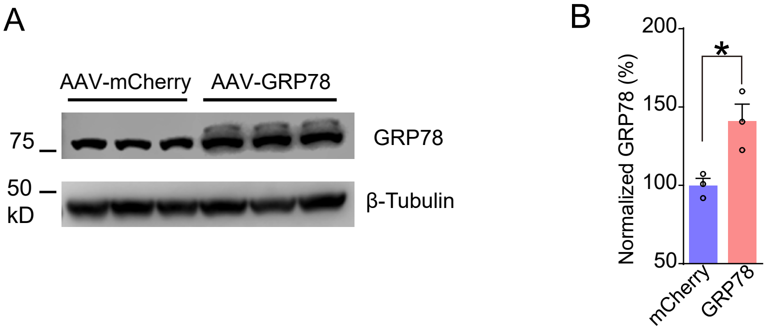

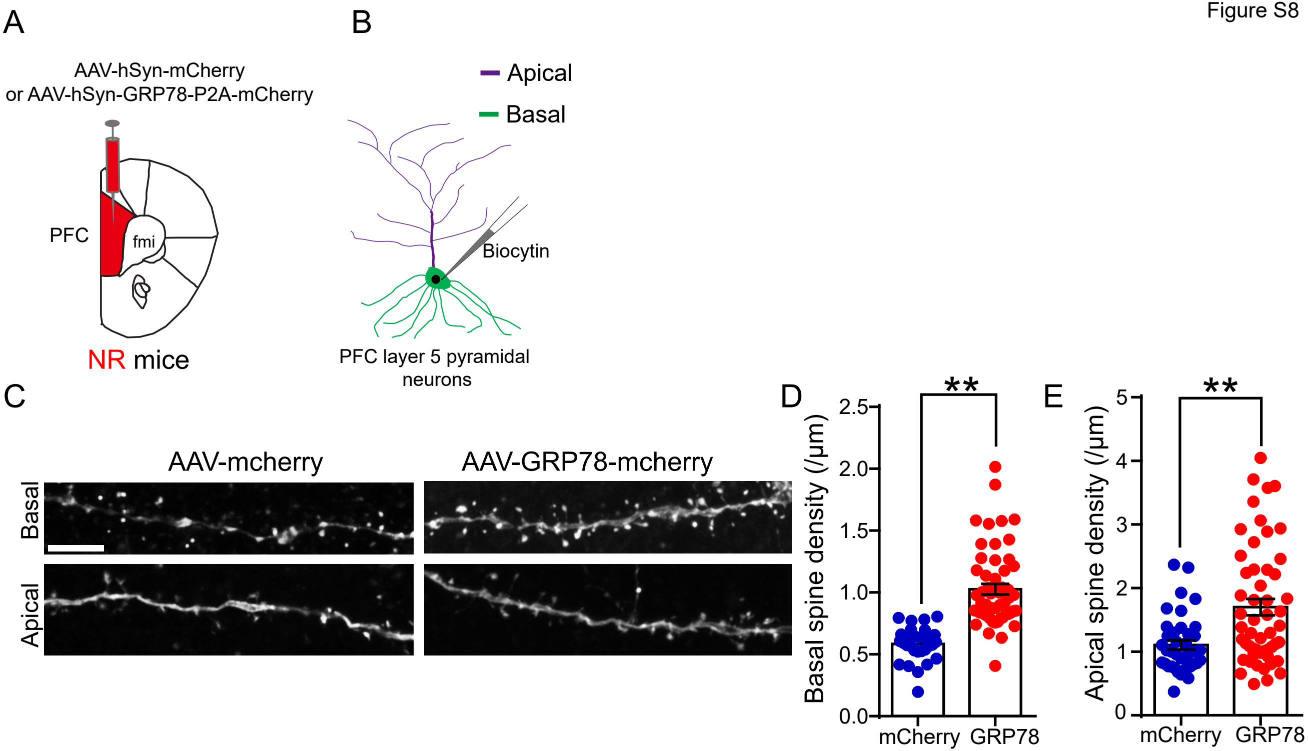

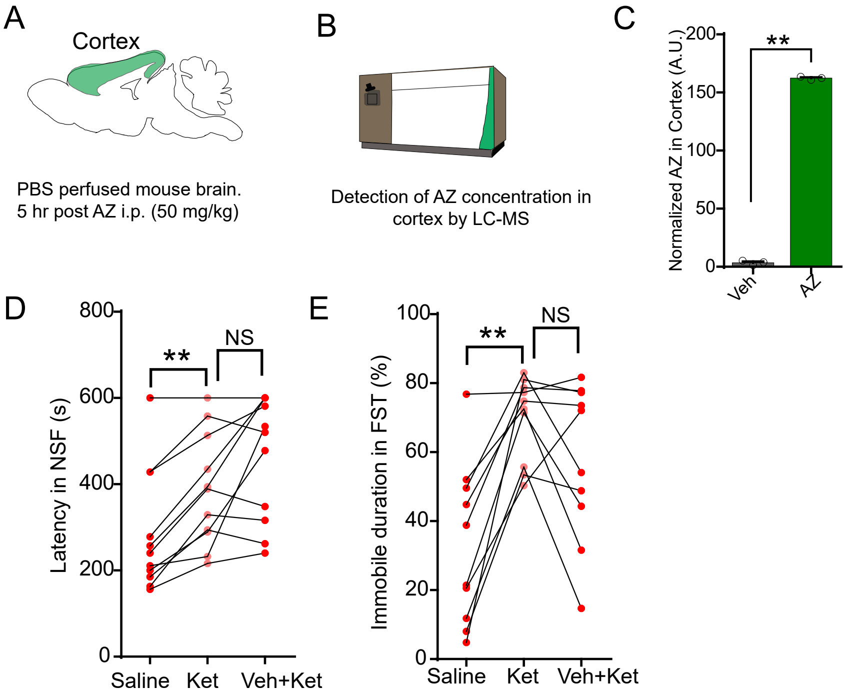

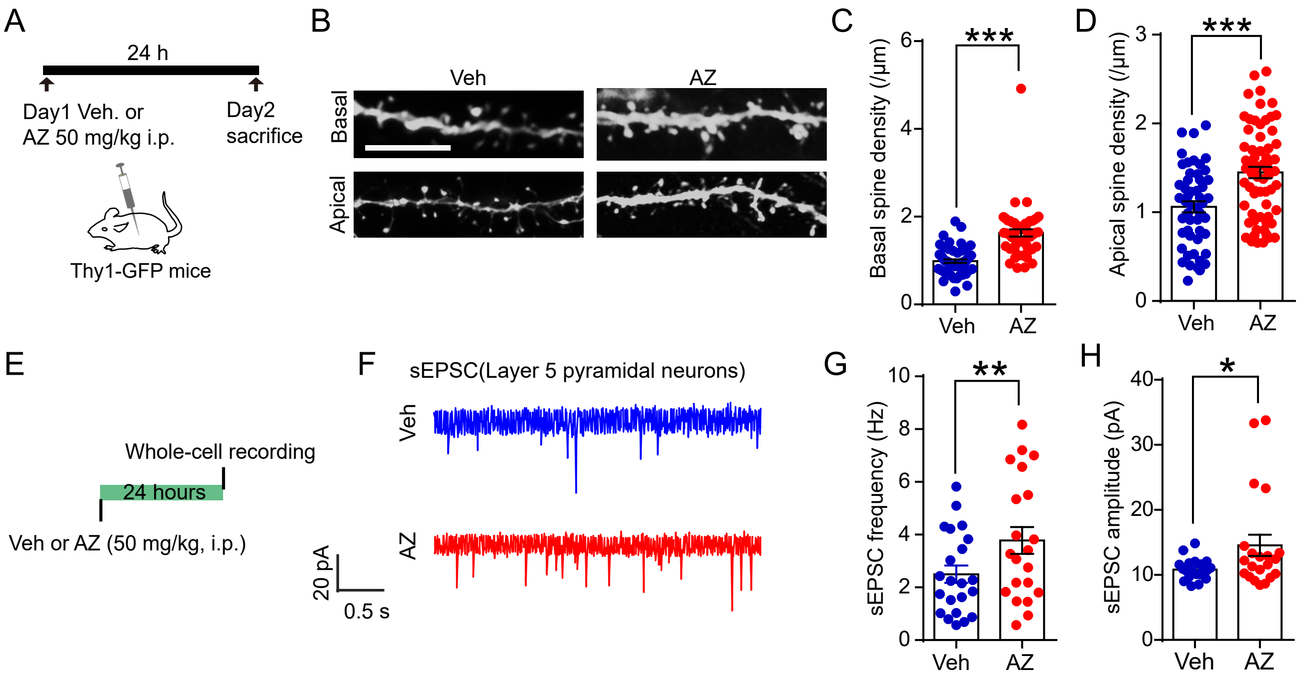
